# Multi-cellular phenotypic dynamics during the progression of breast tumors

**DOI:** 10.1101/2025.04.16.649085

**Authors:** Louise Gsell, Spencer S. Watson, Iva Sutevski, Matteo Massara, Klara Soukup, Alper Eroglu, Jeff E. Mold, Antony Cougnoux, Johanna A. Joyce, Jean Hausser

**Author notes:** Further information and requests for resources and reagents should be directed to and will be fulfilled by the lead contact, Jean Hausser.

## Abstract

In cancer, improving diagnostics and therapeutic interventions can benefit from understanding the cellular and phenotypic heterogeneity of the tumor microenvironment (TME). In recent years, the TME has been profiled at an unprecedented level of detail by performing single-cell RNA sequencing (scRNAseq) on patient samples. However, from patient samples, studying the temporal dynamics of the TME using patient samples has been challenging. Interrogating the temporal dynamics of the TME is critical to understand how inter-tumor heterogeneity is organized into a temporally ordered sequence of causes and consequences in cellular events.

Here we survey the temporal dynamics of the TME by performing longitudinal scRNAseq on mouse breast tumors at different progression time points of tumor progression. We reveal multi-cellular phenotypic dynamics that follow one out of three possible temporal patterns: stable colonization, wave-like, or progressive increase. In particular, IFN-responsive cancer cells, *GzmB* + cytotoxic T cells, as well as C1q macrophages, progressively increase in parallel with tumors progression.

These findings establish the single-cell types and phenotypes in a progressing breast tumor, and determine when these cellular players enter and leave the TME.

## Introduction

Along with cancer cells, tumors are home to a diversity of cell types that define the TME, including immune, stromal, and endothelial cells ^1–4^. Depending on the tumor type, cells of the microenvironment can make up a substantial fraction of the tumor mass ^5,6^. Both the abundance and mutational stability of the non-cancerous cells found in the TME make them an attractive therapeutic target ^1^.

Immune checkpoint blockade is an example of a therapeutic strategy that targets cells in the TME, more specifically T cells ^7^ or Natural Killer (NK) cells ^8^, and can achieve long-term remission in a subset of cancer patients. Other strategies are being developed: vaccines that target dendritic cells (DCs) ^9^, compounds that stimulate anti-tumor immunity by tumor-associated macrophagess (TAMs) ^10–12^, and more ^3^.

Both the abundance and phenotypes of different cell types within the TME determine patient prognosis ^13^. For example, in most cancer types, cytotoxic T cell (Tc) — which can kill cancer cells via effector genes such as granzymes — associate with better prognosis, while regulatory T cells (Tregs) generally indicate worse prognosis ^14,15^. Among TAMs, polarization towards an immunostimulatory/M1-like phenotype usually confers a better prognosis while immune-suppressive/M2-like or C1q phenotypes are associated with worse patient outcome ^13,16,17^. Thus, understanding the cellular and phenotypic ecosystem of the TME could benefit diagnostic accuracy and the development of more-targeted therapeutic interventions.

Beyond histology, single-cell technologies such as flow cytometry, mass cytometry, and scRNAseq have proven very informative to survey the diversity of cell types and phenotypes of the TME ^18–28^. Applying these technologies to patient tumor samples has established atlases of the many cell types of the TME and of their phenotypic heterogeneity^18–20,24–29^. Such studies have revealed that phenotypic heterogeneity can be patient-specific or shared across cancer types, explored how phenotypic heterogeneity associates with genetic, microenvironmental, and cell-autonomous factors, and characterized associations with prognosis ^18–20,29^.

The majority of single-cell TME studies have surveyed the cellular architecture of tumors at one single time-point ^18–23,30^. However, in doing so, the coordinated sequence of cellular and phenotypic events through which the tumor co-opts the microenvironment and progresses is lost. Determining this sequence of events can potentially reveal temporal logic in the inter-tumor variation of the TME. This is similar to how understanding a movie is easier when the scenes are presented in chronological order. In particular, it can narrow down causalities in cellular coordination, based on the principle that cause must precede consequence in time ^31^. It can suggest why different tumors acquire different TMEs by determining branching points in TME trajectories ^24^. Characterizing the time dynamics of the TME could serve to optimally target therapeutic interventions depending on the stage of tumor growth and also suggest mechanisms of resistance to therapy ^32,33^. Thus there is a need to characterize the multi-cellular temporal dynamics of the TME, to serve as a reference for the process of tumor progression, with its single-cell types, their phenotypes and functions, and to determine when these enter and leave the TME.

Surveying the time dynamics of the TME in patients with solid tumors is challenging. Sampling tumors is challenging, due in part to the potential harm-benefit balance for patients and the logistics of clinical practice ^34^. Moreover, repeated biopsy samplings of the tumor could perturb its biology and even alter its fate ^34^. In addition, fractional biopsy samples may not be representative of the whole tumor ^34^. To overcome these obstacles, clinical grade and tumor size can be used as a proxy for time ^32^. But stage and size are still a proxy, and temporal variation is not the only source of patient-to-patient variation: cancer genetics, host genetics, and environmental and physiological factors may also be critical contributing factors ^35–38^.

To address these challenges, inbred mouse cancer models can be used^6,11,12,30,39^. For example, a longitudinal scRNAseq experiment characterized breast tumors at four stages of growth using the MMTV-PyMT genetically-engineered mouse model (GEMM) ^40^. In this study, the authors predominantly focused on the heterogeneity and dynamics of breast cancer stem-like cells. Beyond the dynamics of cancer cells and stromal cell, the phenotypic dynamics of immune cells remain to be interrogated in depth. Exploring this in GEMMs such as the MMTV-PyMT model can be challenging because tumor initiation kinetics are difficult to control and because of the development of multifocal tumors. To address this challenge, allograft cancer models can be used.

In a single-cell multi-organ longitudinal study of an allograft mouse melanoma model, Davidson et al. ^39^ found that cells of the TME and their phenotypes are temporally regulated through interactions that can be identified by intersecting single-cell gene expression data with ligand-receptor databases ^39^. To maximize the depth of the phenotypic survey of the TME, cancer cells were excluded from this previous study. Howerver, cancer cells are central to the architecture, as they respond to and modulate the signaling environment of the TME — interferons, interleukins, TNF, and more — which impacts anti-tumor immunity ^33,41^. Thus, it is of interest to characterize the phenotypic dynamics of cancer cells along with those of other cells in the TME.

To address these questions, we study herein the single-cell transcriptional dynamics of a breast tumor model. In doing so, we survey the whole tumor to obtain a holistic view of the coordination between cell types and phenotypes, including cancer cells. We explore cell types that have rarely been surveyed in the context of animal solid tumor animal models so far, such as innate lymphoid cellss (ILCs) ^42^.

We find that the TME of breast tumors has dynamics that continuously unfold over the four-week time-course of our study. Within the first two weeks post-tumor induction, tumors are stably colonized by monocytes and DCs, while stromal cells disappear from the tumor. This is followed by wave-like dynamics of T helper (Th) cells/Tregs/ILCs characterized by their absence in week 2, peak activation during the third week, and decreased abundance during the fourth week. Finally, interferon (IFN)-responsive cancer cells, *GzmB* + Tc, as well as *C1q* + TAMs increase over the course of the 4-week time-course.

## Results

### A scRNAseq time-course of breast tumor progression identifies tumor-associated cell types and their temporal dynamics

We performed a time-course experiment in an immunocompetent allograft model originally derived from a MMTV-PyMT GEMM of breast cancer (Methods). The MMTV-PyMT model expresses the Polyoma middle T (PyMT) viral antigen under the control of the mammary-specific MMTV promoter ^43^. This GEMM recapitulates the stages of human breast tumor progression, including distant metastases, and expresses biomarkers that define human breast tumor subtypes such as loss of estrogen and progesterone receptors and expression of ErbB2/HER2 ^44^.

Cancer cells were modified to express EGFP and Luciferase to facilitate non-invasive monitoring of tumor growth. Tumors were initiated by injecting 500 cells into the mammary fat pads of C57BL/6J mice. Tumor growth followed a reproducible trajectory across individuals, reaching the 1 cm^3^ volume endpoint in 4 weeks (Figure 1A).

**Figure 1:**
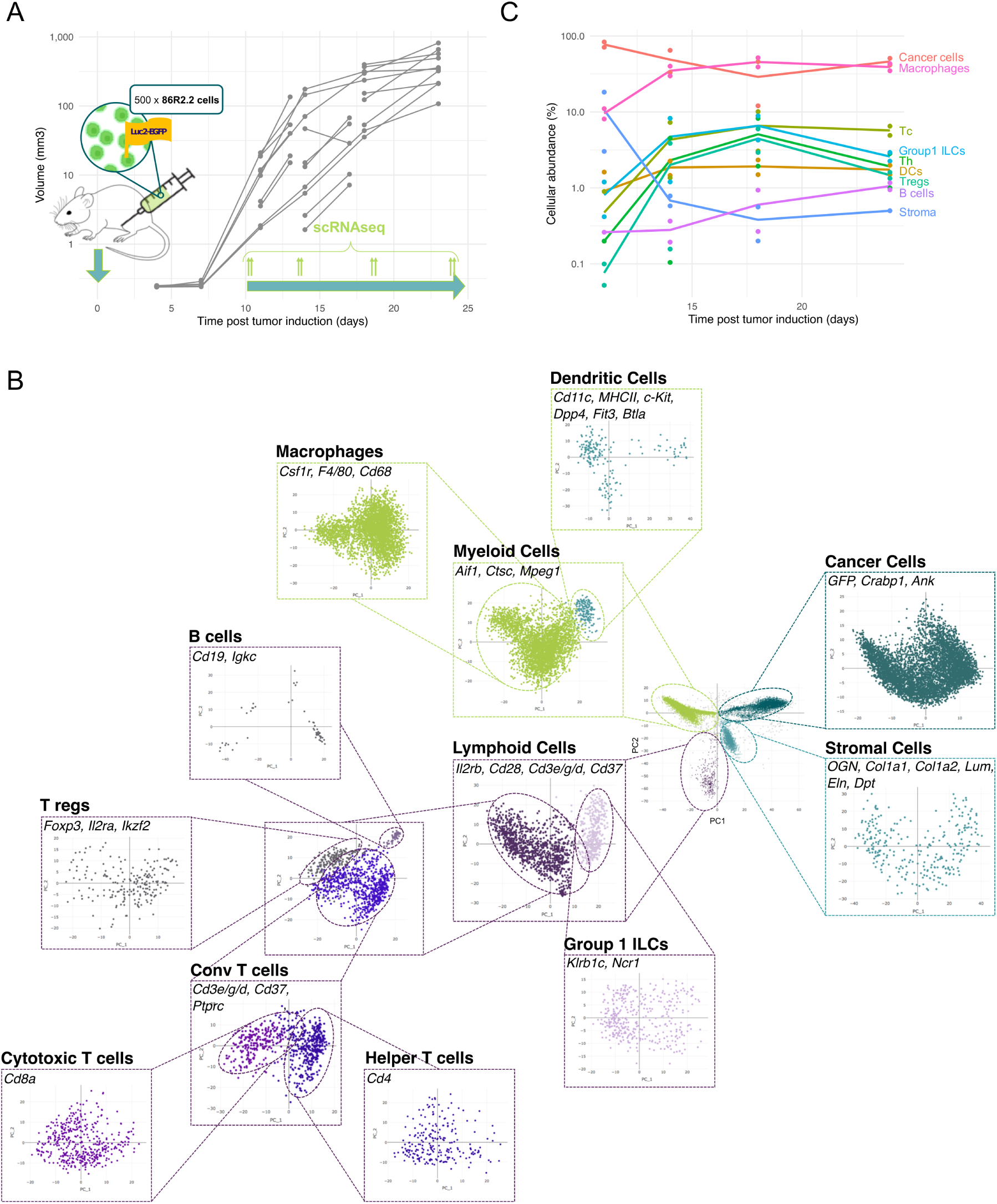
Performing scRNAseq on tumors across multiple growth time points reveals the temporal dynamics of the transcriptional heterogeneity of the TME. **A.** An allograft model of breast cancer was used for the experiment. 86R2.2 cells derived from a MMTV-PyMT genetically engineered mouse model in a C57BL6 background were labeled with Luciferase and GFP as reporters. 500 cells were injected in the mammary fat pads of 30 immuno-competent C57BL6 mice. Tumor growth was monitored non-invasively by bioluminescence imaging. Continuous lines represent longitudinal bioluminescence measurements from the same individual. Tumors were harvested at four time points and scRNAseq performed in duplicates. Y-axis: tumor volume estimated from the bioluminescence signal (Methods). **B**. Hierarchical principal component analysis coupled with Louvain clustering (colors) identifies 9 cell types in this data. Cell type markers expressed in each cluster appear in each box. Axes represent the two first PCs. **C**. The temporal dynamics of the cellular composition of the TME has a biphasic structure. Tumors are initially rich in cancer and stroma, and stroma is subsequently diminished as the immune system colonizes the TME. In a second phase, Th cells, Tregs and group 1 ILCs cells recede while cancer cell abundance increases.

To capture the multi-cellular temporal dynamics of the TME during these 4 weeks, we collected tumors in duplicates, at equally-spaced time points between the day at which tumors were first large enough to be resected and the day at which they reached their volume endpoint (*i.e.* days 11, 14, 18 and 24 post tumor induction, Figure 1A). Tumors were dissociated into single cells and scRNAseq was performed.

State-of-the-art approaches^45,46^ to identify the type of single cells can face challenges in detecting novel cell types and in interpreting the structure and the biology of cellular heterogeneity (Methods). To address these challenges, we used an iterative hierarchical approach to identify cell types and phenotypes. We projected single-cells on their first principal components (PCs) which are weighted linear mixtures of genes that best capture transcriptional heterogeneity. We then looked for low-dimensional structures in the transcriptional heterogeneity of the different cell types: clusters, continua, or geometric shapes.

Principal component analysis (PCA) of single-cell gene expression profiles from all time points and replicates indicated four clusters of cells, which we isolated using Louvain clustering ^47,48^ (Figure 1B). Analysis of cluster-specific genes indicates that these clusters correspond to cancer cells (*GFP*, *Crabp1*, *Ank*), myeloid cells (*Aif1*,*Ctsc*, *Mpeg1*), lymphoid cells (*Il2rb*, *Cd28*, *Cd3e/g/d*, *Cd37*), and stromal cells (*Col1a1/2*, *Lum*, *Eln*, *Dpt*) respectively (Figure 1B). Comparing the gene expression profiles of each cluster with bulk RNAseq profiles from FACS-isolated cell populations confirmed these cell types (Table S1).

To identify the type of each cell more specifically, we iterated PCA and Louvain clustering until cells no longer form separable clusters in the space defined by the first PCs. This procedure identifies nine cell types that express specific markers: cancer cells (*GFP*, *Crabp1*, *Ank*), macrophages and monocytes (*Cd68*, *Csf1r*, *F4/80*), DCs (*Cd11c*, major histocompatibility complex class II (MHCII), *c-Kit*, *Dpp4*, *Flt3*, *Btla*), group 1 ILCs (*Klrb1c/NK1.1*, *Ncr1/NLp46*), Tc (*Cd8*), Th cells (*Cd4*), Tregs (*Foxp3*, *Il2ra*, *Ikzf2*), B cells (*Cd19*, *Igkc*), and stromal cells (*Col1a1*, *Col1a2*) (Figure 1B).

Temporal dynamics differ substantially between cell types (Figure 1C). Initially, at day 11, cancer cells represent 77% of sequenced cells within the tumor, together with stromal cells (11%) and macrophages and monocytes (10%). Within the next two weeks, the stromal component is essentially lost (*<* 1%). The proportion of cancer cells also decreases, while the proportion of innate and adaptive immune cells increases: macrophages and monocytes increase from 10% to 39% while T cells, DCs and Group 1 ILCs increase 3 to 10-fold.

Time dynamics were comparable across replicates (Figure 1C, Figure S1A). To further validate cell type abundances quantified by scRNAseq, we performed hyperplexed immunofluorescence imaging (HIFI)^49^, profiling the abundance of 6 cell types in 11 tumor samples (2-3 sections per sample, Figure S1B,C). Overall, the abundance of cell types as quantified by HIFI matched their abundance as quantified by scRNAseq (Figure S1D, Methods).

Beyond these cellular dynamics, our longitudinal scRNAseq data reveals the phenotypic heterogeneity of each cell type and the temporal dynamics of these phenotypes, which we next investigated.

### The prevalence of IFN-responsive cancer cells, C1q TAM and Tc progressively increases over time

#### Cancer cell heterogeneity is described by proliferation-vs-survival and IFN-response gradients

Cancer cells, the most abundant cell type in our experimental setting, have a transcriptional heterogeneity described by two features.

First, the transcriptomes of cancer cells fall on a curved line in gene expression space (Figure 2A–B). One end of the line represents proliferation, with cells expressing genes involved in mitosis, DNA replication, and cell cycle (Figure 2A–B, Table S2, Figure S2A–B). Cells at the other end of the line express genes involved in glycolysis, the Hypoxia Inducible Factor (HIF) pathway, extracellular matrix (ECM) production, adhesion and organization, and iron uptake (Figure 2B, Table S2) suggesting a hypoxia survival phenotype ^50^. This heterogeneity is found at all time points (Figure S2C).

**Figure 2:**
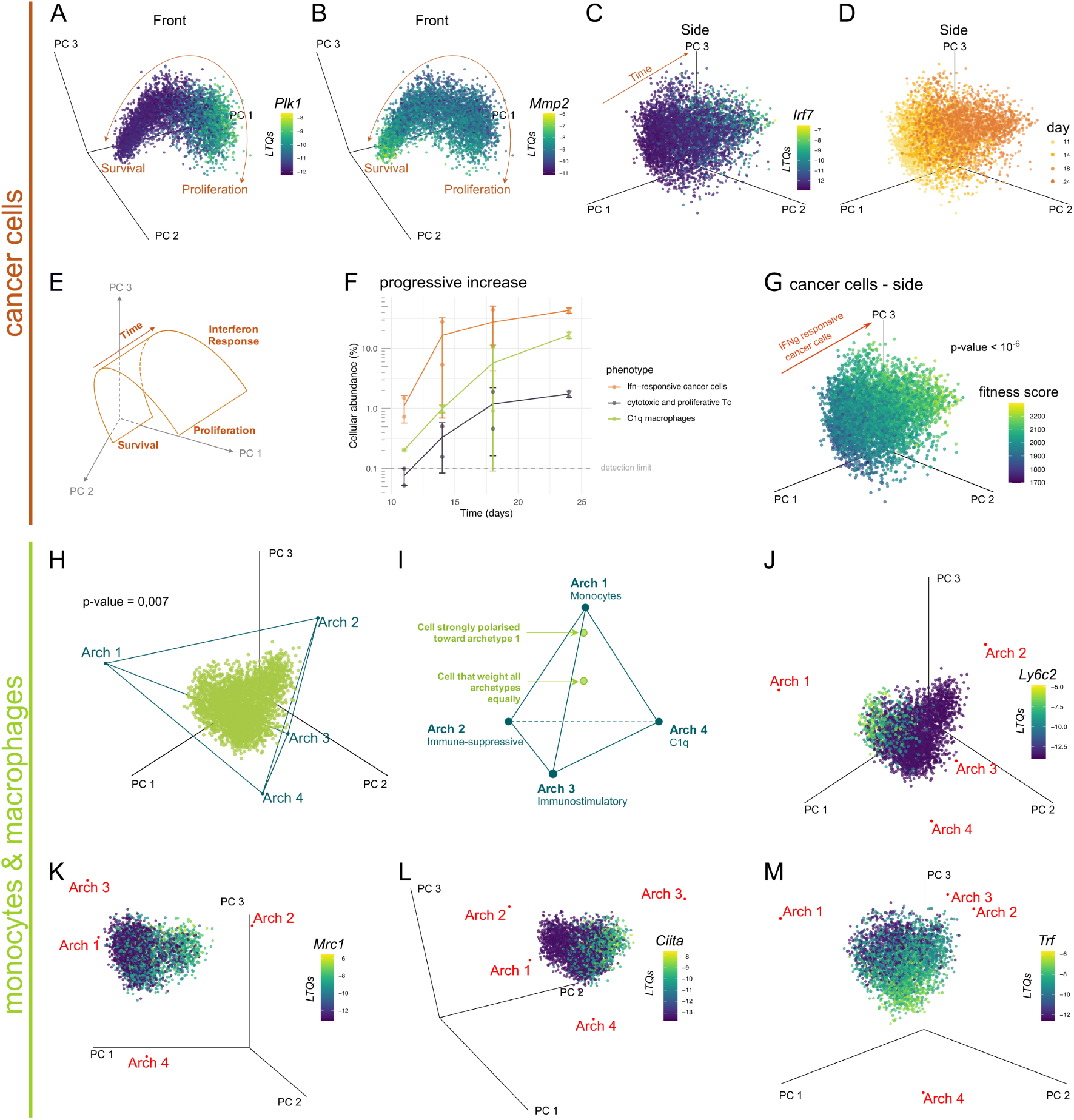
The prevalence of IFN-responsive cancer cells, cytotoxic Tc and C1q macrophages progressively increases along tumor growth. **A-E and G**. The transcriptional heterogeneity of cancer cells describes a linear continuum of proliferation-survival polarization and a gradient in IFN signaling and response. Dots: cancer cells from all time points. Axes: first three PCs of cancer cell gene expression. Color bar: LTQs, a measure of gene expression ^113^. **A**. At one end of the linear continuum, we find high expression of *Plk1*, a cell division gene. **B.** *Mmp2*, an ECM remodeling enzyme associated with tumor invasion, follows the inverse gradient. **C**. Expression of *Irf7* is orthogonal to the survival-proliferation continuum. **D**. Temporal changes in the transcriptional heterogeneity of cancer cells follow the IFN-response gradient. **E**. Summary of the transcriptional heterogeneity of cancer cells. **F**. The abundance of IFN-responsive cancer cells, proliferative Tc and C1q macrophages progressively increases over the four-week experiment. **G**. IFN-responsive cancer cells have highest predicted fitness among cancer cells. Color bar: Cancer cell fitness score computed from the CRISPR experiment of Lawson et al. (2020). P-value: Wilcoxon’s rank sum test on fitness scores of the 5% closest cells to archetype 3 versus other cancer cells (Figure S4B). **H-M**. The transcriptional heterogeneity of macrophages and monocytes is a weighted average of four archetypal phenotypes. Dots: macrophages and monocytes from all time points. Axes: first three PCs of gene expression. **H**. Individual cells fall on a simplex in the first three PCs. P-value: t-ratio test. **I**. Single cell from macrophage / monocyte cluster can be described as a weighted average of the four end-points (archetypes) of the 3D simplex: monocytes, M2-like/immune-suppressive, M1-like/immunostimulatory, C1q macrophages. **J-M**. Cells close to the different archetypes express specific phenotypic markers. **J**. *Ly6c2*, a monocyte marker, is specifically expressed in cells closest to archetype 1. **K**. *Mrc1*, an M2 marker, is expressed in cells closest to archetype 2. **L**. Cells closest to archetype 3 upregulate *Ciita*, part of the M1-like/immunostimulatory macrophage signature. **M**. Cells closest to archetype 4 upregulate *Trf*, an iron uptake protein.

Second, over time, cancer cell heterogeneity progresses along an axis orthogonal to the proliferation-survival continuum (Figure 2C). This time axis (Figure 2D) corresponds to increased expression of genes involved in IFN signaling — IFN regulatory factors (*Irf7*) (Figure 2C), IFN-induced transcripts (*Ifit*, *Rnasel*), IFN receptors (*Ifnar1*, *Ifnar2*, *Ifngr1*) — and effectors of IFN-response: adenosine deaminase acting on RNA (ADAR), antigen processing and presentation genes (*Lgmn*, *Cathepsin S*, *Tap1*, *Tap2*, *Tapbp*, *Cd74*, HLA-DR) (Table S2). There is also increased expression of the EGF/PDGF pathway and components of other signaling pathways including *Pdgfra*, *Map3k1*, *Fos*, *Pik3r1*, *Stat1*, *Stat2*, *Stat3*, *Stat5a* (Table S2).

Thus, the transcriptional heterogeneity of cancer is characterized by a time-invariant proliferation-survival continuum and a time-associated IFN-response phenotype (Figure 2E–F). This progressive increase in the IFN-responsive phenotype could be accessory, induced by changes in the signaling microenvironment of cancer cells, but without evident benefit for the fitness of cancer cells. Alternatively, adopting an IFN-responsive phenotype could confer a fitness benefit to cancer cells so that the progressive increase of this phenotype is the consequence of evolutionary selection. In support of this hypothesis, IFN response has been identified as a determinant of immune escape in multiple studies ^41,51–53^.

To test if this progressive increase of the IFN-responsive phenotype is the product of evolutionary selection *in vivo*, we computed a fitness score for each cancer cell. This score weights the fitness contribution of each gene — as measured by a CRISPR screen aimed at identifying immune escape genes ^41^ — by the expression of that gene. Single cancer cells with a higher ability to escape immunity are thus expected to have higher fitness scores. We find that fitness scores are the highest in cancer cells with an IFN-response phenotype (Figure 2G, *p <* 10*^−^*^6^). This supports the hypothesis that adopting an IFN-responsive phenotype confers a fitness benefit to cancer cells *in vivo*.

The central role of IFNs in immune signaling suggests an immune dimension to the fitness benefit of adopting an IFN-responsive phenotype. To explore the nature of this fitness benefit, we next investigate immune cells whose phenotypic heterogeneity also follows a similar temporal pattern of progressive increase.

#### Macrophage and monocyte heterogeneity polarizes along four fates

As in human breast tumors, macrophages and monocytes are the most abundant immune cells (Figure 1C)^23^. The transcriptional heterogeneity of macrophages and monocytes is poorly captured by phenotypic clusters (Methods). Instead, transcriptional heterogeneity is well described by a continuum shaped as a three-dimensional simplex, a pyramid with a triangular basis (*p* = 0.007 t-ratio test, Figure 2H). The significance of this observation is that any point in a 3D simplex can be uniquely described as the weighted average of the 4 endpoints of the pyramid. These end-points, termed archetypes in the field of unsupervised machine-learning ^54^, can be determined automatically by dedicated algorithms^55,56^.

The biological interpretation of the observation that macrophages and monocytes fall on a simplex is that the transcriptomes of these cells can polarize continuously along four fates — the four archetypes of the pyramid. Heterogeneity arises because individual cells weigh these four archetypes differently (Figure 2I). Consequently, characterizing the heterogeneity of macrophages and monocytes is distilled down to characterizing the four archetypes of the pyramid.

To do so, we analyze the scRNAseq data to identify genes and pathways whose expression is associated with each archetype using the ParTI algorithm ^55^. We supplemented public pathway databases with monocyte and macrophage phenotypic signatures, curated from the literature (Figure S2D) ^57–61^.

The first archetype is associated with markers of activated monocytes, the undifferentiated progenitors of macrophages: *Ly6c2* (Figure 2J), *Ccr2*, *Factor X*, *Cd14* (Table S3). These cells also express genes involved in tissue damage response (*Vegfa*, *Mdk* /midkine *Mdk*) and fibrosis (*Tgfb1*, *Fn1* /fibronectin, *Col1a1* /collagen).

The second archetype is characterized by high expression of genes associated with immune-suppressive macrophages: *Cstd*, *Mrc1*, *Spp1* /osteopontin, *Pf4* (Figure 2K). We also find upregulation of proliferation markers — *Mki67*, *Top2a* (Figure S2E, Table S3). Pathway analysis identifies upregulation of genes involved in wound healing and angiogenesis consistent with the function of immune-suppressive macrophages (Table S3)^62^. A signature of perivascular cells is associated with this polarization, consistent with previous findings (Figure S2F)^63^. Also consistent with a pro-tumoral role of the immune-suppressive state, the cells upregulate *Igf1* (*p <* 10*^−^*^6^, Table S3) which is associated with tumor progression in a diversity of cancer types ^12,64,65^. MHCII expression is low at this archetype, suggesting weak antigen presentation to T cells (Figure S2G).

The third archetype is characterized by genes associated with inflammation and marker genes of immunostimulatory polarization: the MHCII transactivator *Ciita*, pro-inflammatory cytokines *Cxcl9*, *Cxcl10*, *Tnf* and *Txnip* coding for the thioredoxin interacting protein (Figure 2L, Table S3). Pathway analysis reveals upregulation of genes involved in antigen processing and response to IFNs, in agreement with the established pro-inflammatory component of this polarization (Table S3). Consistent with the previously proposed antitumoral role of immunostimulatory TAMs, this phenotype is most abundant in a tumor sample that showed spontaneous regression (Figure S2H).

The fourth archetype expresses marker genes of C1q macrophages ^66^ such as the genes of the C1q complement *C1qa*, *C1qb*, *C1qc* which initiates the complement cascade, *Selenop* and the *Apoe* extracellular vesicle gene (Figure 2M, *p <* 10*^−^*^6^, Table S3), along with genes of the MHCII complex (Figure S2G). While the function of C1q macrophages is less clearly established, they were found to associate with bad prognosis in breast cancer^17^ as well as T cell exhaustion and tolerance ^66^, whereas C1q deficiency is a strong determinant of systemic autoimmunity ^67^. These findings support an anti-immunity effect of C1q macrophages.

The prevalence of the C1q TAMs archetype progressively increases during tumor progression, similar to the IFN-responsive cancer phenotype (Figure 2F).

These parallel dynamics of cancer cells and TAMs suggest crosstalks between these two cell types, in which cancer cells and TAMs influence each other’s transcriptomes. To test for such crosstalk and examine their human relevance, we co-cultured differentiated THP1 cells (dTHP1, a human macrophage model) with MDA-MB-231 cells, a human breast cancer cell line, grown as 3D spheroids (Methods, Figure S3A).

Compared to mono-culture, co-culturing cancer cells with macrophages increased the growth rate and the compactness of spheroids (Figure S3B-C). Gene expression profiling of macrophages upon exposure to cancer cells *in vitro* showed that macrophages up-regulate *C1Q* genes and *APOE* (Figure S3D-F), as in archetype 4 *in vivo*. Among the strongest regulated genes of macrophages, we also observed up-regulation of S100B — a secreted calcium binding protein overexpressed in immune-suppressive TAMs ^68^ associated with poor prognosis in cancer ^69^ — and down-regulation of MHCI (*B2M*, *HLA-A*) and *CLIC1* — a regulator of inflammation in macrophages ^70^ (Table S4).

These observations suggest differentiation of monocytes into C1q macrophages by cancer cells in the human context.

#### Tc heterogeneity is structured along two phenotypic clusters with a shared activation gradient

The transcriptional heterogeneity of Tc cells is described by two clusters (Figure 3A-B).

**Figure 3:**
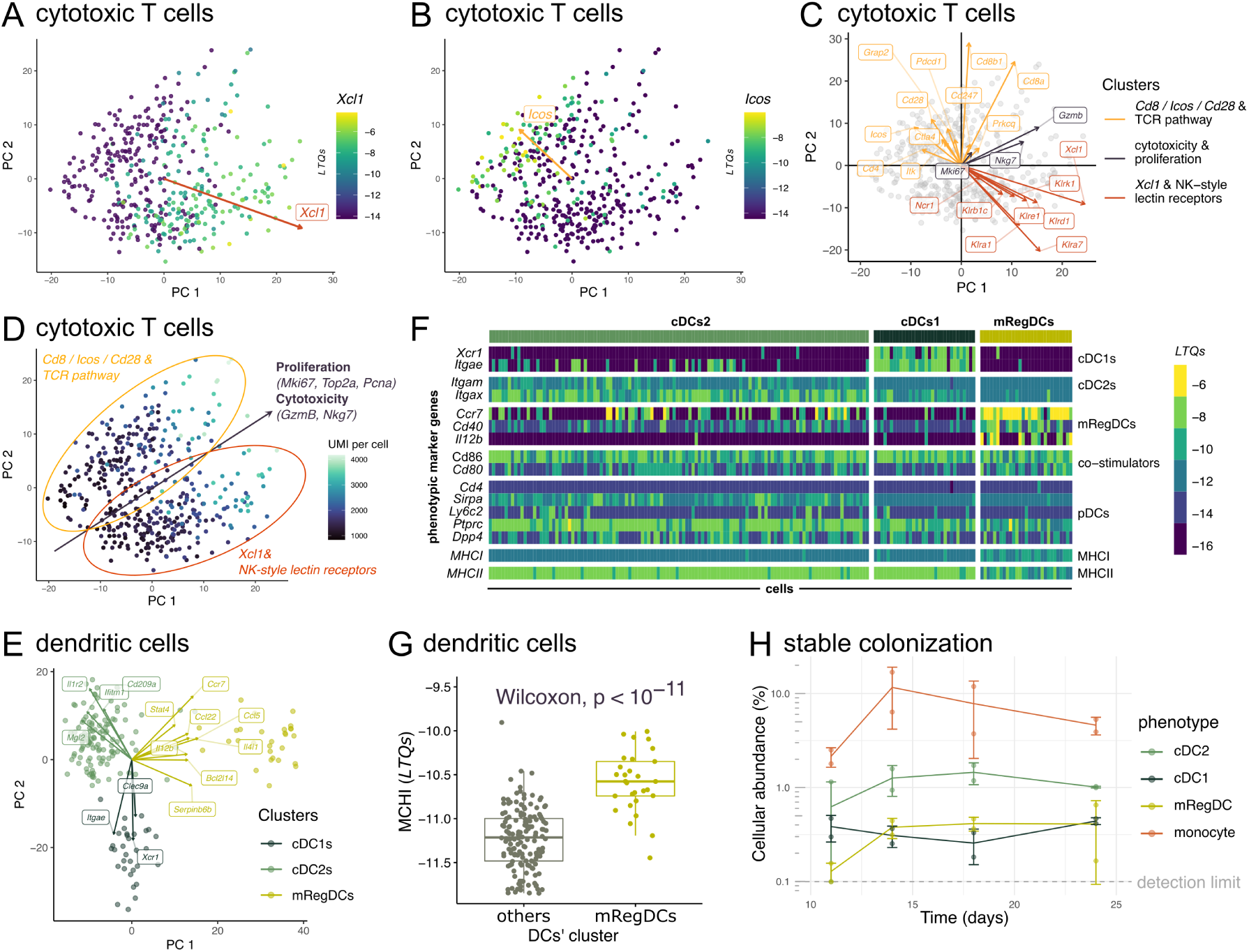
The transcriptional heterogeneity of cytotoxic T cells is characterized by expression of TCR vs NK lectin receptors genes and a cytotoxicity/proliferation gradient while DCs adopt three established phenotypes. **A-E**. Tc cells and DCs form phenotypic clusters. Labels represent pathways significantly associated with PCs. Arrows represent gene expression gradients, with the length of arrows representing the strength of the gradient. **A-D**. Cytotoxic T cells can adopt two phenotypes characterized by expression of TCR or NK lectin receptors genes, with a shared cytotoxicity and proliferation gradient. **A**. Tc cells that score high on PC1 upregulate the *Xcl1* chemotaxis factor which facilitates recruitment of DCs. Color bar: Xcl1 expression (LTQs). **B**. Tc cells that score high on PC2 upregulate the co-stimulatory receptor *Icos*. **C**. Each PC is driven by genes with shared function in TCR or NK lectin receptor signaling. Tc cells that score high on both PCs express genes involved in proliferation, cytotoxicity. **D**. Tc cells can adopt two phenotypes, characterized by expression of *Cd8*, *Icos*, *Cd28* and genes from the TCR pathway or by expression of *Xcl1* and NK-style lectin receptor genes. Both phenotypes share a gradient of activation and transcriptional activity. Color bar: number of UMI per cell, a proxy for transcriptional activity. **E-G**. DCs can adopt three phenotypes: cDCs1, cDC2, and mregDC. **E**. Genes with the strongest expression gradients associate specifically with one DC phenotype. **F**. Expression patterns of DC markers identify the phenotypes of the three DC clusters. Expression of MHCI and MHCII represents the average expression of all genes that comprise the MHC class I and II, respectively. **G**. MregDCs express significantly higher levels of MHCI compared to classical DCs. **H**. The temporal dynamics of mregDC, cDCs1, cDC2 and monocytes follow a stable colonization pattern of the TME. Error bars: one standard deviation.

One cluster upregulated *Xcl1* (Figure 3A), a chemotaxis factor for DC recruitment via the DC-specific *Xcr1* receptor, as well as the Natural Killer (NK)-style lectin receptors *Klrb1c*/*Nk1.1*, *Ncr1*, *Klre1* /*Nkg2I*, *Klra1* /*Ly49A*, *Klrd1* /*Cd94*, *Klra7* /*Lgl-1* /*Ly49g*, *Klrk1* /*Nkg2D* (Figure 3C).

The second cluster upregulates *Cd8a*/*Cd8b*, the co-stimulatory receptors *Cd28* and *Icos* ^71^ (Figure 3B-C), along with other genes of the T cell receptor (TCR) pathway (*p <* 0.003; *Grap2*, *Pdcd1*, *Ctla4*, *Prkcq*, *Cd247*, *Itk*, *Lcp2*, *Cd4*).

Markers of naive/memory vs effector Tc phenotypes^72–75^ do not explain the two transcriptional clusters (Figure S4A, Figure S5). Exhaustion could be another explanatory factor for the two clusters. However, markers of Tc exhaustion previously used in scRNAseq studies like *Havcr2* /*Tim3* and *Tigit* are not detected in our cells (Figure S5), while other exhaustion markers are found in both clusters (*Tox*, *Lag3*, *Pdcd1* /*Pd1*, *Tnfrsf18* - Figure S5, Table S5)^75^. Instead, the data suggest two phenotypes of Tc, polarized towards (1) recruiting DCs and expressing ILCs receptors or (2) expressing co-stimulatory receptors (Figure 3D).

In both clusters, we find a gradient of T cell activation (Figure 3D), with increased expression of the cytotoxic effector *GzmB*, the degranulation marker *Nkg7* ^76^ (Figure 3C), proliferation markers (*Mki67*, *Top2a*, *Pcna*, Figure S4B) and higher unique molecular identifier (UMI) content per cell suggesting a higher transcription activity ^77^.

The prevalence of the cytotoxicity and proliferation phenotype increases progressively over time, thus tracking the dynamics of IFN-responsive cancer cells and C1q TAMs (Figure 2F). This coupling between the IFN-responsive cancer phenotype and a cytotoxicity and proliferation T cell phenotype are consistent with a selection pressure exerted by T cells on cancer cells, and with potential involvement of C1q TAMs.

The transcriptional heterogeneity of Tc shows an expression gradient of the DC chemotactic factor *Xcl1*. This raises the question as to the extent of transcriptional heterogeneity found in DCs, which represents 1–3% of cells in our datasets, depending on the time point (Figure 1C).

### Monocytes and DCs stably colonize the tumor within two weeks post induction

The transcriptional heterogeneity of DCs is characterized by three clusters, each expressing specific genes (Figure 3E). One cluster expresses *Xcr1* — the *Xcl1* receptor — and *Itgae*/*Cd103*. The expression of *Itgae*/*Cd103* in this cluster suggests a classical dendritic cell 1 (cDCs1) phenotype ^78–80^.

A second cluster expresses *Itgam*/*Cd11b* (Figure 3F) highest of all three clusters (*p* = 0.001, Mann-Whitney’s rank sum test), suggesting a classical dendritic cell 2 (cDC2) phenotype ^78–80^. These cells also upregulate *Itgax* /*Cd11c* (Figure 3F) and represent the most abundant DC phenotype in our data set (66%). Other genes are specifically upregulated in these cells — the Il1-receptor *Il1r2*, *Cd209a*/DC-SIGN, *Mgl2*, *Ifitm1* /CD225 — most of which have previously been reported to function in initiating Th cell response ^81–83^.

The data also show a third phenotype. The plasmacytoid dendritic cells (pDC) phenotype can be ruled out based on the undetectable expression of defining pDC markers in these cells (*Itgax*, *Cd4*, *Sirpa*, *Ly6c*), or expression lower than in other clusters (*Ptprc*, *Dpp4*) (Figure 3F). Instead, upregulation of DC maturation genes such as *Ccr7*, *Cd40* and *Il12b* in this cluster suggests a phenotype of mature DCs enriched in immunoregulatory molecules (mregDCs) ^80,84^ (Figure 3F). This phenotype represents a state that cDCs1s and cDC2s can acquire upon sensing a cellular antigen, leading to antigen presentation, immune stimulation and migration ^80,84^. MregDCs were recently identified in lung tumors ^78,84^. Here, we provide evidence for mregDCs in the context of breast tumors.

While cDCs1s were previously proposed to initiate Tc-mediated anti-tumor immunity, we propose that anti-tumor immunity could be driven even more potently by mregDCs. This hypothesis is supported by two observations. First, MHCI expression is enriched in mregDCs compared to cDCs1s and cDC2s (Figure 3F, *p <* 10*^−^*^6^ Figure 3G), suggesting a greater potency in cross-presenting antigens to Tc. Second, while the *Cd86* co-stimulator is found in all clusters, mregDCs show highest expression of the *Cd40* and *Cd80* co-stimulators (Figure 3F, Mann-Whitney test, *p <* 0.0001 and *p* = 0.0001 respectively).

The temporal dynamics of the three DC phenotypes follow a pattern of stable colonization. The prevalence of the phenotypes is initially low at day 11 with the cDCs1, cDC2, and mregDC phenotypes making 0.2-0.6% of cells in the tumor (Figure 3H, Table 1). Subsequently, the prevalence of all three phenotypes stabilizes at 0.4-1%. The cDCs1 phenotype accumulates earlier in the tumor than cDC2s and mregDCs. This is consistent with the previously proposed hypothesis that mregDCs differentiate from cDCs ^84^.

**Table 1:** Summary of the multi-cellular phenotypic dynamics of progressing breast tumors. Phenotypic invariants appear in the first row. Phenotypes specific of a given time-point appear in subsequent rows.

| day | Tc | macrophages and monocytes | cancer | DCs | Th | Tregs | innate lymphoid | Stroma |
| --- | --- | --- | --- | --- | --- | --- | --- | --- |
| **invariant** | two clusters: Xcl1+/NK-like, Cd8/costimulation/TCR | - | proliferation-survival continuum | cDC2, cDC1 (3:1 ratio) | Th1 response gradient | - | mix of NK-ILC1 cells (group 1 innate lymphoid) | - |
| **11** | absent | immune-suppressive macrophages | lfn-responsive and response low | - | absent | absent | low cellular abundance, cAMP response, TGFb | present |
| **14** | low proliferation and cytotoxicity | immune-suppressive macrophages, monocytes | lfn-responsive and response medium | mregDCs | proliferating | proliferation | NK cells activation | absent |
| **18** | intermediate proliferation and cytotoxicity | immunostimulatory macrophages, monocytes | lfn-responsive and response high | mregDCs | matrisome, antigen processing and presentation | activation | ILC1 activation | absent |
| **24** | high proliferation and cytotoxicity | C1q, monocytes | lfn-responsive and response high | mregDCs | decrease in abundance | decrease in activation and abundance | decrease in activation and abundance | absent |

MregDCs express *Il12* (Figure 3G) and *Il15* (Figure S4C), two cytokines that expand and activate group 1 ILCs and CD4+ T cells ^85,86^, which we examine in the next section.

Finally, among TAMs, the prevalence of monocytes follows similar temporal dynamics of stable colonization. From 2% of all cells at day 11, the proportion of monocytes settles between 8% and 12% from day 14 onwards (Figure 3H, Table 1).

### Group 1 ILC, Th, and Treg show a wave-like activation pattern

#### Group 1 ILCs heterogeneity differentiates NK and innate lymphoid cell type 1s (ILC1s) cells, with a gradient of activation

In contrast to other immune cell types — TAMs, T cells, DCs — the transcriptional heterogeneity of ILCs has not been investigated in the context of mouse cancer models to date. Interpreting the phenotypic diversity of tumor-associated ILCs is therefore challenging in the absence of a map of the transcriptional heterogeneity of ILCs.

To address this, we scatter group 1 ILCs on the first two PCs of transcriptional heterogeneity (Figure 4A).

**Figure 4:**
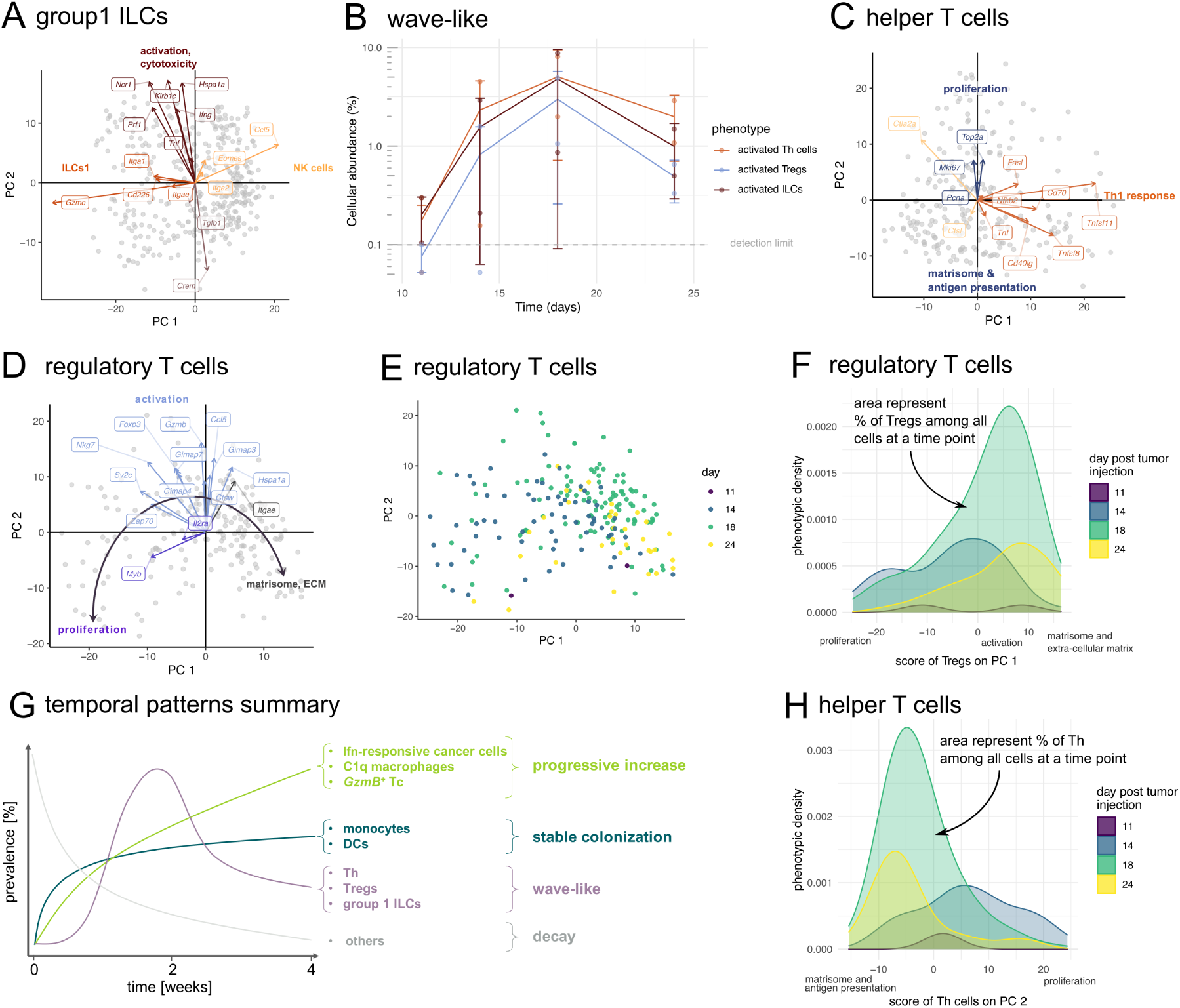
The temporal dynamics of Th cells, Tregs and ILCs are characterized by a wave-like pattern. **A-C**. The transcriptional heterogeneity of Th, Tregs and group 1 ILCs is described by an activation gradient. **A**. Group 1 ILCs are a mix of ILC1s (low PC1) and NK cells (high PC1), with a shared activation / cytotoxicity gradient along PC2. **B**. Th transcriptional heterogeneity is described by a Th1 response gradient along PC1, and an axis of matrisome and antigen presentation versus proliferation along PC2. **C**. A linear phenotypic continuum of proliferation – activation – matrisome, ECM phenotypes characterizes the transcriptional heterogeneity of Tregs. **D**. Time dynamics of Tregs follow PC1. **E**. Treg phenotypic heterogeneity shifts from low PC1 (day 11) to medium PC1 (day 18) to high PC1 (day 24), indicating a temporal shift from a proliferative phenotype initially, to an activated phenotype at intermediate times, to a matrisome and extracellular phenotype at the latest time points. The phenotypic density (y-axis) represents the prevalence of Tregs with a specific phenotype (defined as PC1 score), so that the area under each curve equals the fraction of Tregs at this time point. **F**. During tumor progression, the transcriptional heterogeneity of Th cells shifts from a proliferative phenotype to a matrisome and anti-gen presentation phenotype. **G**. The temporal dynamics of activated Th cells, Tregs and ILCs follow a wave-like pattern. **H**. The multi-cellular phenotypic dynamics of the TME during the progression of a breast tumor model is summarized by three temporal patterns: progressive increase, stable colonization and wave-like.

We find that PC1 associates with NK cells versus ILC1s phenotype. Cells that score high on PC1 express markers found in NK cells and not in ILC1s ^87^ — the *Ccl5* /*Rantes* cytokine which recruits multiple immune cell types to tumors, *Itga2* /*Cd49b*, *Eomes*/*Tbr2* (Figure 4A) — as well as genes involved in the matrisome and in collagen formation (FGSEA *p* = 0.02 and 0.007 respectively, Table S6). In contrast, cells that score low on PC1 express ILC1s-marker genes — the cytotoxic agent *Gzmc*, *Itgae*/*Cd103*, *Itga1* /*Cd49a*, *Cd226* (Figure 4A) — as well as other genes involved in cytotoxicity and the ribosome (Table S6). Cellular UMI counts decrease along PC1, suggesting higher transcriptional activity in ILC1s compared to NK cells (Figure S4D).

The second principal component captures a gradient of activation and cytotoxicity. Cells that score high on PC2 express genes that promote cytotoxicity: the cell lysis factor *Prf1*, the death factor *Tnf*, the inflammatory cytokine *Ifng*, the *Hspa1a* survival factor and the innate lymphoid receptors *Ncr1* and *Klrb1c*/*NK1.1* (Figure 4A). Consistent with PC2 capturing cytotoxicity and activation, cells that score low on this axis express the cAMP-responsive element modulator *Crem* — cAMP has an inhibitory effect in NK cells ^64^ — and the immunosuppressive signal *Tgfb1* (Figure 4A).

The phenotypic dynamics of group 1 ILCs follow a wave-like temporal pattern (Figure 4B). Group 1 ILCs have low abundance at day 11, with cells displaying a cAMP-responsive and immuno-suppressive phenotype (Table 1, Figure S4E). By day 14, we see an increased abundance of NK cells with an activated, cytotoxic phenotype, potentially in response to Il12/Il15 signaling by mregDCs from day 11. The cytotoxic phenotype is then adopted by ILC1s cells at day 18 (Figure S4E). At day 24, ILCs group 1 abundance generally decreases, with a smaller proportion of cells in the activated, cytotoxic phenotype (Table 1, Figure S4E).

#### The heterogeneity of Th cells and Tregs follows a gradient of activation

We finalize our analysis of phenotypic heterogeneity and of its temporal dynamics by turning our attention to Th cells and Tregs.

Although some Th2/Th17 markers are detected in Th cells (Figure S4F) and not all Th1 markers are detected, the transcriptional profile of Th cells fits most closely with a Th1 fate. In addition to the Th1 signature, we detect “immunosuppressive” markers *Pdcd1* /*PD1*, *Lag3*, *Ctla4* (Figure S4F).

To explore the transcriptional heterogeneity of Th cells, we represent single cells on the 2D plane defined by the first two PCs of the data. We find that PC1 represents a gradient of Th1 effector response (Figure 4C). Cells that score high on PC1 express the CD40 ligand (*Cd40lg*) which is induced upon MHCII-associated antigen presentation to Th cells ^86^. These cells express signals employed by Th1 to induce apoptosis: the tumor necrosis factor (TNF), the TNF super-family members *Tnfsf8* and *Tnfsf11*, and the *Fas* ligand (*FasL*). Expression of genes that promote survival such as *Cd70* — a co-stimulatory ligand — and the *Nfkb2* transcription factor also follow PC1, as well as other genes of the TNF-receptor non-canonical NF*κ*B pathway (FGSEA *p* = 0.016), perhaps facilitating the survival of Th cells in the pro-apoptotic signaling environment they establish ^88^. In contrast, cells that score low on PC1 express the cathepsin L protease (*Ctsl*) — which plays a role in antigen presentation and elastin degradation — and its inhibitor *Ctla2a*.

PC2 captures a gradient of proliferation. Cells that score high on PC2 express genes involved in proliferation (*p <* 0.001) while cells that score low on this axis express genes involved in the matrisome (*p <* 0.001) and in antigen processing and presentation (FGSEA *p* = 0.001, Table S7).

The transcriptome of Tregs stands out with respect to conventional T cells (Figure 1B): we find that Tregs do not simply cluster as a subset of CD4+ T cells, as is typically the case in flow cytometry analyses or when clustering T cells based on canonical T cell markers. Tregs express several markers of naive T cells — *Lef1*, *S1pr1*, *Cxcr4* (Figure S6A)^18,75^. The naive phenotype of Tregs could be due to the specific-pathogen-free environment in which the animals are kept, with little infections ^89^. Alternatively, these markers could also indicate memory Tregs as they were previously associate to memory. There is ubiquitous expression of *Maf* — a transcription factor required to differentiate into the *RORγt*+ and T follicular regulatory T cell phenotypes^90^ — the *Icos* co-stimulatory receptor, the *Ctla4* inhibitory receptor, and the *Ikzf2* /*Helios* activation marker.

Representing Tregs on the first two PCs of the data suggests a linear differentiation trajectory (Figure 4D–E). Consistent with a linear differentiation trajectory with a temporal ordering in gene expression, the expression gradients of genes responsible for Tregs heterogeneity are oriented along a continuous arc (Figure 4D). This contrasts with the four directions of gene expression gradients found in group 1 ILCs for example (Figure 4A).

On one extreme of the differentiation trajectory, scoring low on PC1, we find cells expressing the *Myb* cell-cycle driver, the *Il2* receptor *Il2ra* and proliferation genes (*p <* 0.001, Table S8).

In the middle of the trajectory, scoring high on PC2, we find genes involved in T cell activation such as the TCR-associated signal transduction protein *Zap70* and the *Ccl5* /*Rantes* pro-inflammatory cytokine. These cells express higher levels of the Tregs transcription factor *Foxp3* and of the Il2-induced *Cathepsin W* ^91^. In addition, these cells express the *Sv2c* vesicle-associated protein, *Nkg7* which is required for the degranulation of cytotoxic vesicles and the cytotoxic effector *GzmB*. There is also upregulation of the *Gimap4* apoptotic effector and other genes of this family (*Gimap7*, *Gimap3*), as well as *Hspa1a*, proposed to counter the effect of apoptotic signals.

At the end of the trajectory, scoring high on PC1, we find cells expressing the *Itgae*/*Cd103* integrin (Figure 4D) and genes involved in the matrisome, extra-cellular matrix organization and collagen formation (*p <* 0.001, Table S8).

The observed cytotoxicity transcriptional gradient (*GzmB*, *Nkg7*) may be surprising given the immunosuppressive function of Tregs. A mislabeling of Tc as Tregs can be ruled out (Figure S6B). Instead, Tregs have previously been proposed to inhibit Tc and NK cells using cytotoxic gene products ^92^.

The phenotypic dynamics of Th cells and Tregs unfold progressively from day 11 through to day 24, following a wave-like temporal pattern: absence or low abundance at day 11, recruitment and activation at day 14 and 18, before the progressive loss of function and abundance at day 24 (Table 1).

Regarding Tregs for example, time dynamics mainly happen along PC1 (Figure 4E). Tregs are detectable by scRNAseq from day 11 (Figure 4E, Figure 4F, Table 1). At day 14, Tregs score lowest on PC1 compared to other time points, indicating a proliferation phenotype. By day 18, the density of cells with an intermediate and high score on PC1 increases, indicating a shift towards the activated and matrisome/ECM phenotypes. At day 24, the shift towards the matrisome/ECM phenotype continues and Tregs decrease in abundance (Figure 4F, Table 1).

For Th cells, time dynamics occur mainly along the PC2 axis (Figure S6A). Th cells are initially absent at day 11 (Figure 4E, Table 1). At day 14, cells score highest on PC2 compared to other time points, indicating a proliferating phenotype. At day 18, most cells score low on PC2, indicating a shift towards a matrisome and antigen presentation phenotype. Finally, by day 24, Th cells decrease in abundance. In addition, the distribution of Th cells along the PC1 axis of Th1 response was comparable across time points (Figure S6C): this suggests a time-invariant Th1 response gradient, perhaps additionally associated with a spatial component.

## Discussion

The development of therapies aimed at the TME can benefit from understanding its cellular and phenotypic architecture. By studying a mouse model of breast cancer, we find that the TME undergoes continuous changes over four weeks as breast tumors progress (Figure 4G). Tumors initially consist mostly of cancer cells and stromal cells. During the first two weeks of tumor progression, the prevalence of stromal cells decreases while monocytes, cDCs1s, cDC2s, and mregDCs stably establish themselves in the TME. Subsequently, the abundance and activity of Th, Tregs and ILCs follow a wave-like pattern: undetectable initially, their activity peaks during week 3 before receding again during week 4. Finally, IFN-responsive cancer cells, C1q macrophages, and activated *GzmB* + Tc gradually increase as the tumor progresses.

This dataset can serve as a reference for how cancer cells co-opt the tumor microenvironment during tumor progression, as a result of our longitudinal, multi-cell-type characterization. In addition, we sequenced at sufficient depths to profile a sizable fraction of the cells’ transcriptomes (in the order of 100k unique mRNAs), with 22356 Unique Molecular Identifiers (UMIs) per cell on average. Sequencing at this depth places the current dataset on this high end in terms of sequencing depth, thus providing a reference dataset for the deep single-cell transcriptome of progressing breast tumors (Figure S7B).

For the sake of advancing breast cancer therapy, it is important to explore how the cellular dynamics of our mouse model recapitulate those of human breast tumors. Making inter-species comparisons is challenging because phenotypic markers differ across species. For example, *F4/80* /*Emr1* /*Adgre1* is a marker of the mononuclear phagocyte lineage in mice, but in humans, *ADGRE1* is an eosinophil marker. Similarly, *Ly6c* is a mouse monocyte marker but is not expressed in human monocytes. Such inter-species differences in cellular phenotypes, molecular signals and more generally cell biology may explain why mouse treatments have often failed to translate to humans ^93^. Yet, mouse tumors can be compared to human tumors using omics tumor profiling and their dynamics compared using grade or size as proxy of tumor progression. We performed such comparisons using publicly available human tumor data. We found that cellular and gene expression dynamics in mice overlap with inter-sample variation of human breast tumors from day 14 of our experiment (supplemental note 1). Beyond this direct comparison, there is scientific and medical value in establishing cellular dynamics in mouse tumors because some of the tumor biology of mouse models does translate to humans. For example, targeting immune checkpoint inhibitors was first demonstrated in mice ^94^.

The present data can inspire future mechanistic and clinical studies. For example, our analysis suggests that adopting an IFN-responsive phenotype confers a fitness benefit to cancer cells in vivo. This could inspire therapeutic strategies aimed at blocking IFN-responsive genes in cancer cells. Similarly, the progressive increase of C1q macrophages suggests reprogramming of macrophages by cancer cells, a hypothesis that we now tested in human experimentally (Figure S3). Finally, the mregDC phenotype was previously found to limit antitumor immunity in non-small cell lung cancer (NSCLC)^95^, which inspired therapeutic strategies aimed by mregDCs: blocking IL-4 signaling, combining immune checkpoint inhibitors with dendritic cell therapy based on Flt3L and *α*CD40 ^95^. Finding mregDCs in our breast tumor model suggests the same approaches for breast cancer therapy.

The observed cellular and phenotypic time dynamics suggest hypotheses about the underlying microenvironmental mechanisms of tumor progression. For example, the dynamics of stable colonization observed in cDCs1s, cDC2s, mregDCs, and monocytes calls for a mechanism to explain the temporally stable prevalence of these cell types. One possibility is that the progressing tumor establishes different niches with specific cellular signatures — for example, a niche dominated by cancer cells, another niche consisting mainly of dendritic cells and monocytes — with these niches expanding at the same rate as the tumor grows, thus preserving cellular proportions.

The hypothesis that niches expand at the same rate conflicts with the progressive increase in the prevalence of IFN-responsive cancer cells, C1q macrophages, and activated *GzmB* + Tc during tumor progression. These phenotypes remain distinct from a steady state, much like the tumor which grows steadily and never reaches a stable size in our experiment (Figure 1C). To explain this, one possible hypothesis is that IFN-responsive cancer cells, C1q macrophages, and activated *GzmB* + Tc localize in a specific immune escape niche which expands faster than other niches during tumor progression. In support of this hypothesis, IFN-response by cancer cells has been shown to induce classical and non-classical MHCI expression to down-regulate ILC and T cell anti-tumor immunity respectively ^96^. An alternative hypothesis is that constant generation of neo-antigens, fueled by genomic instability of cancer cells, drives a sustained increase in the rate of *GzmB* + Tc recruitment, which leads to the selection of the IFN-responsive cancer phenotype at the expense of other cancer phenotypes, perhaps with the support of C1q macrophages. Indeed, cancer cells with an IFN-responsive phenotype express genes associated with increased fitness in the face of cytotoxic selection by Tc ^41,51–53,97–99^. Niche-independent hypotheses are also possible: DC recruitment could be proportional to the overall number of cancer cells, with selection favoring the IFN-responsive cancer cell phenotype over time.

Further hypotheses are needed to explain the wave-like pattern in Th cells, Tregs, and ILCs activity and its timing, with a peak during week 3. Dynamical systems theory suggests that the time scale of immune dynamics is set by the time needed for the rates of immune cell recruitment, proliferation, and differentiation to stabilize in the tissue that originally triggered the immune response, and balanced by the rate of immune cell regression and death in that tissue ^100^. Upon acute viral or bacterial infection, for example, the wave of adaptive immunity typically unfolds over two weeks: following an innate immune response in the first hours to days of infection, antigen detection and amplification of T and B cells requires approximately one week, with another week necessary for the infection to resolve ^101–103^. Since cancer cells also introduce antigens in the form of foreign proteins — and in this model system, additionally the PyMT oncogene, GFP, and luciferase — one could expect similar dynamics ^104^. Yet, instead of peaking during week 1 and resolving during week 2, Th, Tregs and ILCs activity unfold progressively from day 11 through to day 24, with a peak in week 3 (Table 1). It would be interesting to investigate what causes the two-week delay in Th cells, Tregs, and ILCs activation and why this activation wave is not sustained in tumors. One hypothesis could be that a critical mass of antigenpresenting cells (DCs, macrophages) is needed to initiate Th cells/Tregs/ILCs response, which delays their activation. Th cells/Tregs/ILCs activation may lead to the death of cancer cells lacking the IFN-response phenotype and, subsequently, resolution of the Th cells/Tregs/ILCs response. Future studies could investigate this hypothesis by interfering with Th cells/Tregs/ILCs response or engineering a constitutive IFN-response phenotype in cancer cells.

Previously identified ligand-receptor communication networks could also explain some of the phenotypic dynamics. For example, Tc are characterized by a gradient of expression of the *Xcl1* ligand. Increase in *Xcl1* + Tc precedes *Xcr1* + mregDCs, consistent with recruitment of mregDCs by *Xcl1* + Tc ^86^. MregDCs are in turn characterized by expression of *Il12* (Figure 3F) and *Il15* (Figure S4C), two cytokines that expand and activate group 1 ILCs (alongside T cells) ^85^. Accordingly, we observe an increase of activated group 1 ILCs from day 14, following the detection of mregDCs at day 11.

We propose the here-identified sequence of events as an overall pattern of multicellular dynamics in progressing breast tumors. Yet, inter-individual comparison of tumors cellular composition suggests that tumor may run on individual-specific clocks: for example, the 631-1 scRNAseq sample from day 14 is most similar to the 629-5 from day 18, rather than the 636-4 replicate sample of day 14 (Figure S1A, Figure S8A). Expanding the analysis by HIFI, we find a similar axis of variation as in scRNAseq opposing lymphoid cells to cancer cells, with important individual-specific dynamics (Figure S8B). For example, certain tumors sampled at day 11 feature important lymphoid infiltrates expected of more advanced tumors (Figure S8C-E). This is notable given that animals have the same age, genotype, and identical tumor genetics. The origin of this inter-individual variation is still to be determined: inter-individual variability is not significantly associated with factors such as volume, individual weight, tumor growth rate or tumor size (Figure S8A,B,F). Future studies can explore the origin of individual-specific tumor progression speeds, perhaps in terms of factors such as blood metabolic profiles, group social dynamics within a cage, individual activity, immune repertoire, individual stochasticity or stochastic incidence of metastases which could modulate the primary tumor progression rate. One challenge in resolving factors driving individual microenvironmental dynamics will be to identify the type of single cells from tissue sections with high specificity: in our analyses, the limited size of the panel as well as technical difficulties resulted in many cells whose identity could not be determined specifically, and whose prevalence varied strongly across samples. This limits our ability to describe precise cellular temporal dynamics and identify factors driving inter-individual variability in these dynamics.

The transcriptional heterogeneity of macrophages and monocytes is well described by the geometry of a three-dimensional simplex. This observation generalizes the two-pole model (M1-M2) of macrophage heterogeneity^57,105,106^ to four poles: monocyte, immunostimulatory/M1-like, immune-suppressive/M2-like, and C1q. In addition, the three-dimensional simplex geometry suggests an evolutionary interpretation of the phenotypic heterogeneity of the transcriptomes of single macrophages and monocytes: macrophages and monocytes may need to perform the four functions implemented by the four above-mentioned archetypal phenotypes but cannot perform all functions at once due to trade-offs between these functions ^107,108^. Under this hypothesis, a theorem ^107,108^ predicts that optimal single-cell transcriptomes should fall on a simplex, as is observed here.

The transcriptional heterogeneity of DCs shows a clear pattern of three clusters, with two conventional DC phenotypes and one mregDC phenotype. MregDCs were previously reported in the context of human and mouse lung tumors ^78,84^. Here we provide evidence for their presence in mouse breast tumors. We also explore the transcriptional heterogeneity of group 1 ILCs, a cell type for which there is limited single-cell data in the context of cancer. Here, group 1 ILCs are found in the form of ILC1s and NK cells, and follow a similar activation program, with NK cell activation preceding activation of ILC1s by several days. In conclusion, our study establishes a temporal sequence of the progressing breast TME, with a single-cell repertoire of its cell types, their phenotypes, and functions, and when these cells enter and leave a multi-step tumor progression cascade. Future research should explore the intercellular communication signals and selective pressure responsible for the observed temporal patterns.

### Limitations of the study

Compared to the allograft model we used, spontaneous mouse cancer models present the advantage of better mimicking the early stage of the tumor. However, mimicking early tumorigenesis was not our primary goal: instead, we aimed to characterize the temporal cellular dynamics of a progressing tumor. For this sake, there are strong reasons to prefer an allograft model: (i) the time point of tumor initiation is precisely controlled, which is essential to position tumor samples on a time axis, (ii) confounding factors such as anatomical site and multi-foci tumors can be controlled, (iii) cancer cells can be labeled to verify that tumors follow representative and reproducible growth dynamics through precise, non-invasive, longitudinal monitoring of tumor growth, as well as specifically identify cancer cells by scRNAseq. For these reasons, we intentionally chose a breast allograft over a spontaneous model. In addition, we addressed one of the most important drawbacks of allograft models by titrating the cells down to the minimum required for tumor initiation. Indeed, most studies using PyMT-derived allografts initiate tumors with hundreds of thousands of cells. Instead, initiating our tumors with 500 cells positions our model closer to human tumors which begin from a small number (one) of cells. Departing from the standard practice to initiate allograft tumors with hundreds of thousands of cells may potentially lead to different immune cell recruitment kinetics compared to previous allograft studies. The relevance of our allograft study to human breast tumors is supported by the finding that cellular and gene dynamics in mice overlap with inter-sample variation of human breast tumors from day 14 of our experiment (supplemental note 1).

Studying a single cell line could offer a biased picture of the micro-environmental dynamics of mouse allograft tumor models. To address this, one could have compared these dynamics across allograft models, at the risk of merely establishing an atlas of cell-line specific artifacts. Instead, we intentionally focused on deeply characterizing a tumor model that had the most human disease-relevant features (Lin et al. ^44^, supplemental note 1).

We study here a model of a hot, immunogenic tumor which succeeds in escaping the immune system despite ample recruitment of immune cells into the tumor. Such tumors are expected to have different immune-related features compared to less immunogenic tumors. The immune features of the present allograft tumor model overlap with those of human tumors, an observation that supports its relevance (supplemental note 1).

## STAR Methods

### Cell line

86R2.2 cell cultures were initially derived from a MMTV-PyMT GEMM in a C57BL/6J background ^6^ and cultured in DMEM supplemented with 10% fetal bovine serum and 1% penicillin/streptomycin. Cells were transfected with GFP and Luciferase for non-invasive monitoring of tumor growth by a lentiviral vector encoded in a pFUGWPol2-ffLuc2-eGFP plasmid ^109^. Following transfection, cells were passaged 5 times to remove lentiviral particles. GFP-positive cells were selected by FAC-sorting.

### Animal and Tumor model

C57BL/6J mice were housed and bred in the conventional animal facility of Centre des Laboratoires d’Epalinges, Lausanne, Switzerland in individually ventilated cages with environmental enrichment and ad *libitum* food and water. All animal studies were first approved by the Institutional Animal Care and Use Committees of the University of Lausanne and Canton Vaud, Switzerland under the license VD3072. Mice were injected at 6-10 weeks of age in the fourth left mammary fat pad with 500 86R2.2 cells in a solution 1:1 with Matrigel (Corning) using a 26G needle.

Mice were closely monitored on a daily basis to assess weight changes, level of activity, and the presence of any discomfort or pain. No adverse effects (weight loss, seizure development, poor grooming, labored breathing) were observed, in agreement with our previous experience with this cancer model.

### Caliper and BLI measurements

Tumor growth was monitored weekly by bioluminescence imaging and caliper measurements to verify the reproducibility of tumor growth dynamics across individual animals and to ensure that tumors did not grow beyond the maximum size according to humane endpoint conditions (1 cm^3^).

For bioluminescence imaging (BLI), mice were anesthetized using 3% isofluorane. Luciferin (Promega) was pre-heated at 37 degrees Celsius and injected intraperitoneally at a concentration of 10 mg/mL and a dose of 150 mg/kg. Prior to imaging, isofluorane was reduced to 2% to avoid depressing respiration. Mice were arranged in order according to their individual identifier and eye gel was applied to prevent eye desiccation. Using the IVIS system (Perkin-Elmer), a calibration image was taken with a 5 sec exposure time at a pixel binning of 4 to estimate luminescence. Starting 5 min post luciferin injection, an image was taken every 2.5-5 min until 15-30 min post-injection, long enough to ensure that photon flux peaked. Imaging time (5-90 seconds) and pixel binning (1-16) were set to collect 20’000 photon events in a single exposure. Following imaging, tumor size was quantified by caliper, and the volume estimated as *V* = ^1^ *πL* × *l*^2^ mice were weighted and positioned on a warm surface to recover from anesthesia. After the mice fully recovered from anesthesia, they were returned to the cage.

The BLI images were analyzed using Living Image (Perkin-Elmer) and custom R scripts to determine the temporal dynamics of the photon flux following luciferin injection, summed over the region of the tumor. Maximal photon flux was used as a correlate of tumor size, as previously reported ^110,111^.

### Tumor tissue preparation

We collected tumor samples on days 11, 14, 18, and 24 post-tumor initiation (Figure 1A). At each time point, tumor size and growth was assessed the day before so as to select two tumors whose volume and growth rate were average compared to tumors at that time point. Typical tumor volume and growth rate were established using 10 animals whose tumors we monitored non-invasively by bioluminescence imaging along the whole course of the experiment.

Animals were sacrificed by lethal anesthesia with an intraperitoneal injection of pentobarbital (150 mg/kg). Adequate depth of anesthesia was confirmed by observing loss of reflex and absence of change in respiratory rate associated with manipulation and/or firm ear, toe, or tail pinch. Abdominal skin was disinfected by ethanol. Following confirmation that a suitable anesthetic plane was attained, the animal was positioned in a supine position, and a midline sternotomy was made with a fresh, sterile scalpel. The heart was visualized, and the left ventricle was identified. A 25-27 gauge needle attached to a syringe containing 10 mL of PBS was carefully inserted through the myocardium and into the left ventricle, and the solution was infused. Simultaneously, the right atrial appendage was localized and a small incision was made with sharp scissors in order to accommodate the additional volume being administered and to evacuate the vascular tree. These procedures were carried out as quickly, but as carefully, as possible while the heart was still beating to ensure adequate distribution of the PBS to all tissues.

Tumors were localized in the fourth mammary fat pad by palpation and visual inspection. A fluorescent binocular was used to confirm the tumor location on day 11. Following resection, tumors were measured by caliper (width, height) and weighted.

Using a scalpel, 10-250 mg of tumor tissue was cut for single-cell RNA sequencing. The tissue was further cut into 1-2 mm-sided cubes. The cubes were positioned on a 40 *µ*m filter and washed with 15 mL Hanks’ Balanced Salt Solution (HBSS). Tumor fragments were collected into C-tubes (Miltenyi, Switzerland) using a pipette tip, and 4.7 mL of DMEM Glutamax w/o FBS were poured over the reversed filter to detach any small remaining fragments into the C-tube. Enzymes H (200 *µ*L), A (25 *µ*L), and R (100 *µ*L) from the GentleMACS tumor dissociation kit (Miltenyi, Switzerland) were added to the C-tube. C-tubes were processed using the human tissue dissociation kit program 1 for soft tissues on a GentleMACS OctoDissociator instrument (Miltenyi, Switzerland) for 65 minutes.

The dissociated tumor tissue was filtered through a 40 *µ*m filter into a 50 mL falcon by pre-washing with 10 mL of HBSS, pouring the dissociated tumor tissue on the filter, using a syringe plunger on the filter to gently squeeze tissue through the filter, and washing the filter with 5 mL HBSS to yield 20 mL volume of dissociated tissue.

Dissociated cells were pelleted at 300xG for 10 min at 4 degrees Celsius. The supernatant was discarded, and pellets were re-suspended in 10 mL 1x red blood cell lysis buffer and incubated at room temperature for 10 min. Cells were pelleted again at 400xG for 10min at 4 degrees Celsius, re-suspended in 5 mL of filtered DMEM w/FBS, and counted twice on a Countess automatic cell counter (ThermoFisher Scientific). Viability was quantified by Tryptan Blue staining. Cell counts and viability were estimated by averaging the two measurements.

For single-cell RNA sequencing, a concentration of 1–1.3×10^6^ cells per mL and a volume of at least 25 *µ*L is necessary. To prepare a cell suspension fitting these specifications, cells were pelleted at 300xG for 5 min at 4 degrees Celsius, re-suspended at a concentration of 2600 cells per *µ*L, and filtered through a 35 *µ*m filter, a step which halved cell concentration in our hands. Following filtrating, cell concentration and viability were measured twice again to confirm a concentration of 1–1.3×10^6^ cells per *µ*L. Samples were put on ice and taken to the EPFL Gene Expression Core Profiling facility.

### scRNAseq

Cells were checked for the absence of significant doublets or aggregates and loaded into a Chromium Single Cell Controller (10x Genomics, Pleasanton, CA) in a chip together with beads, master mix reagents (containing RT enzyme and poly-dt RT primers), and oil to generate single-cell-containing droplets. Single-cell Gene Expression libraries were then prepared using Chromium Single Cell 3’ Library & Gel Bead Kit v3.0 (PN-1000075) following the manufacturer’s instruction (protocol CG000183 Rev C). Quality control of the libraries was performed with a TapeStation 4200 (Agilent) and QuBit dsDNA high-sensitivity assay (Thermo Fisher Scientific) following manufacturer instructions. With this procedure, the cDNAs from distinct droplets harbor a distinct and unique 10x cell barcode. The pooled sequencing libraries were loaded onto Illumina NextSeq 500 Flow Cells and sequenced using read lengths of 28 nt for read1 and 132 nt for read2, at a depth of ca 100k reads/cell. The Cell Ranger Single Cell Software Suite v3.0.1 (https://support.10xgenomics.com/single-cell-gene-expression/software/pipelines/latest/what-is-cell-ranger) was used to perform sample demultiplexing, barcode processing, and 3’ gene counting using 10X Genomics custom annotation of mouse genome assembly mm10, to which GFP-luciferase was added (nt 3402-7500 of Addgene plasmid 71394).

CellRanger-filtered UMI count matrices were loaded into Seurat version 4.1.1 ^47^. Genes detected in less than 5 cells and cells with less than 100 detected genes were removed from the data, following common practice. Inspecting the distribution of the percentage of mitochondrial transcripts per cell showed a tail of outlier cells with more than 12.5% of mitochondrial transcripts, which we excluded from further analysis ^112^. The UMI count matrix was then log-normalized using Sampling Noise based Inference of Transcription ActivitY (Sanity) ^113^, an algorithm that uses Bayesian inference to filter out Poisson noise from scRNAseq UMI count matrix. Sanity returns log transcription quotients (LTQs) and associated error bars for each gene.

### Dimension reduction and clustering

To interpret the scRNAseq data in terms of cell types and phenotypes, the state-of-the-art approach consists of grouping cells into clusters of cells with similar transcriptomes ^47^. Clusters are identified in terms of cell types — cancer, stroma, T cells, B cells, and so on — (semi-)automatically by comparison to reference datasets or using a database of marker genes ^114–116^. Resolving cells of each type into phenotypes can be achieved by increasing the granularity of clusters ^78^. This approach is convenient, efficient, and generalizes across experimental settings, in health and disease.

However, the automatized clustering approach to identify cell types and phenotypes has limitations. Cells of types that only partially fit the reference data, or novel cell types missing from the reference can be misinterpreted. The clustering approach to phenotyping can miss structures that may better describe phenotypic heterogeneity than clusters such as continua or geometric shapes. To identify such continua structures among thousands of cells and thousands of genes, cells can be projected on low-dimensional manifolds using methods such as UMAP ^117^ and t-SNE ^118^. But the non-linear character of the axes of variation identified by these methods makes it challenging to interpret the structure and the biology of cellular heterogeneity. Projections are often limited to two dimensions, potentially masking phenotypic structures in three or more dimensions.

To address this, we first determine cell types. We perform a centered unscaled PCA analysis on LTQs of all samples (all time-points and mice) using ade4 R package (version 1.7.19) ^119^. Analyzing all samples together is more efficient as cell type calling is performed only once on the whole dataset, and combining multiple samples increases the density of cells in gene expression space, which facilitates cell type identification.

Scattering single cells on the first three PCs of the gene expression data showed four cell clusters. Cells were assigned to these four clusters using Louvain clustering implemented in Seurat (version 4.1.1)^47^ with the default *k* = 20 neighbors. Louvain clustering has a resolution parameter that determines the granularity of communities: the smaller the resolution, the smaller the identified communities, and thus the higher the number of these communities. Setting the resolution to 0.02 automatically captured the four clusters. To identify the cell type of each cluster, we then determined genes highly expressed and specific to each cluster to look for marker genes of known cell types (Figure 1B). We also compared the average transcriptional profile of each cluster with databases of transcriptional profiles from FACS-isolated cell populations (BioGPS: GNF1M, MOE430)^120^. This procedure identified our four clusters as myeloid cells, cancer cells, lymphoid cells, and stromal cells.

The same process (PCA and Louvain clustering) was then applied recursively to each cluster until cells formed a continuum in PC space and Louvain clustering could not distinguish further clusters without substantially increase the resolution parameter (Figure 1B). This procedure identified eleven clusters. Among those eleven clusters, we find nine cell types: cancer cells, macrophages/monocytes, DCs, B cells, Tc, Th cells, Tregs, ILCs, stromal cells. The remaining three clusters are subtypes of DCs: cDCs1, cDC2, mregDCs (Figure 1B). The resolution parameter was increased to 0.03 when analyzing clusters of cells more homogeneous than the general cellular populations, in order to distinguish cellular subtypes. Among conventional T cells, a resolution of 0.07 was needed to distinguish Th cells from Tc. Similarly, visual examination of DCs suggested three clusters, which were automatically captured by Louvain clustering at a resolution of 0.1.

Stromal cells were few and difficult to assign cell types to so we refrained from determining more detailed cell types. Neutrophils, endothelial cells, and adipocytes were not identified, possibly due to experimental procedures (size-based filtering, enzymatic tissue dissociation, red blood cell lysing buffer).

At each hierarchical level of the PCA and clustering analysis, PCA and clustering were performed only on genes with high signal-to-noise ratio (SNR): these were genes whose signal (standard deviation in expression across cells, as quantified by Sanity) was sufficiently large relative compared to the noise (computed from the average sampling error on LTQs per gene, as estimated by Sanity). We used a cut-off of SNR*>* 0.25 because it performed best at distinguishing the cell types expected in this tumor. Cells with low RNA content were difficult to assign to specific cell types or subtypes. These cells were, therefore, removed prior to clustering, by finding the lower tail of the UMI content per cell. The number of PCs was determined using the elbow criteria (Figure S9)^119^.

In addition, we removed cells in regions of gene expression space with low cellular density and thus uncertain cluster specificity. To do so, we computed the average distance *d* to the nearest neighbors of each cell: *d* is a measure of cellular isolation. Excluding cells with *d* beyond some cut-off can potentially remove cells with uncertain cluster specificity. However, since different clusters have different numbers of cells and thus densities, a unique cut-off on *d* cannot fit all cell types. To address this, we note that, at constant cellular density in gene expression space, *d*^3^ ∝ 1*/n*, with *n* the number of cells in the cluster. Based on visual inspection of clusters in PCs space, we thus set a cut-off on *nd*^3^ to exclude cells with uncertain cluster assignment.

Note that identifying high SNR genes, removing cells with low RNA content and cells falling in regions of low density had to be repeated at each hierarchical level of the PCA and clustering analysis because exclusion criteria are cell-type specific. For example, a cancer cell with low RNA content may have more RNA than a T cell with high RNA content. Thus, eliminating cells with low RNA content was done in a cell-type-specific fashion.

### Interpreting phenotypic heterogeneity

No correction for time-point or sample-effects was necessary to identify cell types (Figure 1B). Within cell types, there was substantial phenotypic overlap across time points (Figure 2D, Figure 4E–F, H, Figure S6C-E). These observations suggest that batch effects are minimal in our experiment.

In addition, we verified that cellular phenotypes were reproducible across duplicates of a given time-point: representing cells in their low-dimensional projection, we verified that cells from duplicates of a given time point had a similar distribution in their low-dimensional gene expression space.

#### Gene expression gradient analysis

Interpreting the phenotypic heterogeneity within each cell type is challenging because scRNAseq profiles the expression of thousands of genes, too many to be considered one by one.

We addressed this using two approaches: (a) interpreting genes with the strongest expression gradients in terms of their function from the literature, and (b) systematic functional enrichment using pathway databases.

To interpret genes with the strongest expression gradients in terms of their function from the literature, we determined the 30 genes with an expression gradient most associated with the first PCs, a number low enough to allow gaining an overview of the literature for individual genes in all cell types, and visualizing these genes on a figure. Genes with the strongest expression gradients were identified according to the sum of squares of their loadings on the first PCs. We searched these genes in the literature to identify their known function in the context of the cell type of interest, with an emphasis on identifying groups of genes that collectively contribute to interpretable phenotypes. Genes of unclear function were discarded for visualization purposes. The complete list of genes and their loadings is reported in section Supplemental Tables. Gene expression gradients were visualized using distance biplots ^119^: genes were projected on the first PCs and their expression gradient was represented as a vector whose length represents the weight of the gene on each PC. Vector lengths were up-scaled a 100-fold for visualization purposes, following common practice in multivariate statistics ^119^ (Figure 3A–C, E, Figure 4A, C–D).

Marker gene and literature search focused on the 30 genes with the strongest expression gradient. This approach risks missing global patterns of heterogeneity. To address this, we interpreted each PC by performing gene set enrichment analysis (GSEA) on the molecular signatures database (MSigDB version 3.0) which offers a catalog of genes involved in many cellular functions and pathways ^121^. GSEA analysis looks for groups of genes that collectively associate with each PC, either positively or negatively. The analysis was performed using fgsea (version 1.21.1) ^122^.

#### Interpreting the simplex geometry of macrophage/monocyte heterogeneity

For macrophages and monocytes, we take advantage of the observation that the phenotypic heterogeneity of these cells is constrained by a 3D simplex. Interpreting heterogeneity in the context of simplex structures is best done by characterizing the phenotypes of the end-points of the simplex. To do so, we use the ParTI Matlab software package ^55^ which was developed for this purpose.

Visual examination of the transcriptional heterogeneity of macrophages and monocytes suggests that this heterogeneity is constrained by a 3D simplex. We test the statistical significance of fitting a 3D simplex to the transcriptional heterogeneity of macrophages and monocytes using the t-ratio test implemented in the ParTI Matlab package ^55^. Briefly, given a high-dimensional dataset such as a gene expression count matrix, ParTI performs PCA and then determines the position of the end-points of the simplex in PC space using algorithms from machine learning and satellite image analysis ^54,56^. These end-points, also known as archetypes in machine learning, define a simplex that encompasses the data. ParTI also quantifies how well the simplex fits the data, and whether such a fit is expected by chance. To do so, ParTI computes the *t*-ratio, defined as the ratio between the volume of the simplex and the volume of the convex hull of the data ^107^. To test if the simplex fits the data better than a random dataset, ParTI computes the *t*-ratio on 1000 shuffled datasets that preserve the distribution of the PC’s loadings but not their correlation. A one-sided *p*-value representing the fitted simplex’s statistical significance is then computed from the *t*-ratio distribution. We find a p-value of *p* = 0.007 (t-ratio test), supporting the hypothesis that a 3D simplex is a good statistical description of the transcriptional heterogeneity of macrophages and monocytes (Figure 2H).

To help interpret the archetypes, ParTI also returns the list of pathways and genes whose expression peaks in cells closest to each archetype, with associated *p*-values (see the previous section), which we used to interpret the phenotypic heterogeneity of the macrophage/monocyte cluster. We tested if adding additional archetypes (5 or 6 instead of 4 archetypes) highlighted genes and pathways not found in the first 4 archetypes. We found that adding a 5th or 6th archetype identified genes and pathways mostly found in the first four archetypes. For this reason, the final analysis includes four archetypes.

We also took advantage of the ParTI methodology to find genes and pathways whose expression peaks at the end-points of the phenotypic surface described by cancer cells (Figure S2A–B).

The transcriptional heterogeneity of TAMs and monocytes is not uniform throughout the simplex: certain transcriptional profiles are more common than others (Figure S2I– J). The observation that the density of the transcriptional profile is not uniform suggests an alternative hypothesis to the simplex structure: transcriptional heterogeneity could be interpreted as 4 clusters that blend into each other due to measurement error.

Several observations are inconsistent with the cluster interpretation. If heterogeneity was explained by discrete phenotypes, there should be groups of macrophages separated by empty space in the principal component analysis. The data conflict with this expectation (Figure 2H, Figure S2I–J).

Phenotypic clusters may still be reconciled with the data under the hypothesis that measurement error is large so that single cells fill in the empty space in between cluster centroids. The data conflicts with this hypothesis for two reasons. First, if the continuum was explained by clusters with measurement error, each vertex should have its own density peak. Yet, there are no density peaks close to the phagocytic TAMs and monocytes end-points (Figure S2I–J). Second, under the four-cluster hypothesis, the width of the density peak should match the measurement error of scRNAseq, as estimated by Sanity. We find that the measurement error is much smaller than the width of density peaks (ellipses on Figure S2I–J).

Thus, the data support a continuum of transcriptional heterogeneity shaped as a simplex better than transcriptional clusters.

### Visualizing phenotypic time dynamics

Visualizing the temporal dynamics of transcriptional heterogeneity of the different cell types is challenging because, except for DCs, phenotypic heterogeneity has mostly a continuous nature in our experiment.

The simplest approach to tracking phenotypic dynamics is to project cells on PC space, and color them by time point (see e.g. Figure 2D). However, this approach suffers from three limitations. (1) Different numbers of cells were profiled in different samples and time points. Time points with a lot of cells can wrongly convey the impression that a new phenotype is appearing, even if this phenotype has the same prevalence in all samples. (2) Changes in the prevalence of phenotypes are difficult to distinguish from changes in the prevalence of cells of this cell type. (3) The eye is more sensitive to changes in abundant phenotypes than changes in rare phenotypes.

For cell types whose phenotypic heterogeneity is well described by a one-dimensional continuum — Tregs, Th cells — we can represent the density of phenotypes at the different time points over their 1D axis of heterogeneity. To allow comparing phenotypic dynamics across samples where different numbers of cells were sequenced, densities are normalized so that the area under the curve represents the prevalence of the cell type at each time point (Figure 4F, H). This normalization allows visualizing phenotypic dynamics, aggregating the effect of cellular differentiation, proliferation death, and migration of cells into and out of the tumor.

This approach generalizes poorly to cell types where multiple axes are needed to capture phenotypic heterogeneity because representing changes in densities over multi-dimensional spaces is not visually compelling. We thus discretize phenotypes and represent how their prevalence changes over time (Figure 2F, Figure 3H, Figure 4B). To compare phenotypic dynamics across samples where different numbers of cells were sequenced, the prevalence of phenotypes was normalized to the total number of cells sequenced at each time point. One drawback of this approach is that it focuses phenotypic dynamics on average effects as opposed to tracking changes in the distribution of phenotypes. Yet, we find discretization necessary to visualize the dynamics of multi-dimensional phenotypes. The approach has the benefit of facilitating the quantification of experimental variability across replicates.

Discretization of cellular phenotypes was guided by gene expression gradient and pathway enrichment analyses (see the previous section).

Macrophages and monocytes were discretized into five clusters, based on the distance to each archetype. First, we identified the 25% closest cells to each archetype and assigned them a phenotype depending on the archetype: C1q, immunostimulatory/M1-like, immune-suppressive/M2-like, and monocyte. The 20% cells closest to the origin were assigned a generalist phenotype, in line with the Pareto optimal interpretation of the simplex structure ^55,107^.

Tc heterogeneity was found to be organized along three categories: *Cd8* /*Icos*/*Cd28* and TCR pathway, cytotoxic and proliferation, *Xcl1* and NK-style lectin receptors (Figure 3C). Each category is associated with a specific region in PC space. In addition, the time axis was found to follow the same direction then the cytotoxic and proliferation gradient (Figure S6D), we thus chose to discretize according to this gradient: cells with positive PC1 and PC2 were labeled as cytotoxic and proliferative Tc (Figure 3C).

In Tregs, Th cells and group 1 ILCs, heterogeneity was captured by two PCs. Of these two PCs, only one was associated with time. The time-association PC was used to discretize these cells, depending on whether cells scored positively or negatively on this PC. PC1 was used to discretize Tregs into proliferating vs matrisome and extra-cellular matrix phenotypes. PC2 was used to discretize Th cells into proliferating vs matrisome and antigen-presenting phenotypes (PC1 captured a gradient of Th1 response that was largely conserved across time points). PC2 was used to discretize group 1 ILCs into activated vs non-activated cells (PC1 separated NK cells from ILC1s).

Cancer cells were discretized according to the expression of *H2-K1* which strongly associates with the IFN-responsive phenotype. H2-K1 expression is bimodal and stratifies cancer cells into *H2-K1* -high and *H2-K1* -low populations. Inspection of the histogram suggests that setting a threshold to LTQs = −7.7 best discriminates the two sub-populations.

### Computing fitness scores of individual cancer cells from CRISPR screening data

Our analysis of cancer cell heterogeneity revealed a time-associated IFN-response gradient. Temporal changes in the prevalence of this phenotype could be explained by changes in the signaling microenvironment or be the result of evolutionary selection *in vivo*. Indeed, IFN-response was shown to play a critical role in cytotoxic T lymphocyte (CTL) evasion in multiple genetic screens ^41,51,52^.

To test the hypothesis that the IFN-responsive phenotype confers a selective advantage to cancer cells in escaping immunity, we scored the fitness of each cancer cell based on its transcriptome, by weighting the fitness contribution of each gene by its expression in individual cells. The fitness contribution of each gene was measured in a CRISPR screen by Lawson et al. ^41^. Lawson et al. ^41^ performed genome-wide CRISPR screens across multiple mouse cancer cell lines, among which two breast carcinoma models: EMT6 and 4T1. Cancer cells were co-cultured with CTLs to induce a CTLs evasion selection pressure. By performing CRISPR screens, they computed the change in the abundance of cells with different KO genes over the course of the co-culture experiment to quantify the selective advantage of each gene as a normalized Z score (NormZ, ‘end’ time-point). To estimate the fitness of single cancer cells, we weighted the NormZ scores from the EMT6 cell lines by the expression of each gene in our data: fitness score = LTQs · NormZ scores.

Representing the fitness score in the space of the first three PCs showed a gradient along the IFN-response phenotype, suggesting that IFN-responsive cancer cells have the highest fitness among cancer cells (Figure 2G). Comparing the fitness scores of the 5% cells closest to the IFN-response phenotype (Figure S2B) vs the remaining cancer cells showed a statistically significant difference (*p <* 10*^−^*^6^, Wilcoxon’s rank sum test). The same observations hold when using the fitness measurements from 4T1 CRISPR screen (Figure S6F).

### THP1 differentiation

THP1 is an acute monocytic leukemia cell line obtained from the American Type Culture Collection. Following treatment with 50ng/mL of phorbol 12-myristate 13-acetate (PMA) these cells become adherent and reproducibly acquire a macrophage-like phenotype, with numerous similarities to primary human macrophages. One million THP1 cells per 75cm2 flask were incubated with PMA for 48h. The cells were then washed twice with PBS to remove the non-adherent cells and the medium was replaced for 24h with cell maintenance medium: DMEM with 10% FBS, Penicillin-Streptomycin, and Mycoexpert. To make single cell suspension the culture medium is removed, the cells washed with PBS, and incubated for 5 minutes with 0.25% Trypsin at 37°C. The protease is quenched by the addition of a medium prior to the evaluation of cell viability. Viability was estimated using a Cellcounter2 and 0.4% trypan blue solution according to manufacturer instructions.

### Spheroid mono- and co-culture

MDA-MB-231 and THP1 cells were tested for mycoplasma and grown in a maintenance medium. All the lines were grown at 37 degrees, 5% CO2, and 100% humidity. No experiment used cells maintained for more than 10 passages to reduce the risk of genetic drift. The MDA-MB-231 (clone A from our lab) and differentiated THP1 cells were washed twice with PBS to remove the non-adherent or dead cells, trypsinized for 5 minutes at 37 degrees, and quenched with maintenance culture medium. If the viability was below 90%, cells were centrifuged at 500g for 5 minutes and the pellet was resuspended in medium. For the spheroid monoculture, 5000 live cells were seeded on 1% agar-coated 96-well plates with round bottom. Reproducible aggregate-like spheroids form in 5 days for the MDA-MB-231 monoculture. For the spheroids co-culture 5000 live MDA-MB-231 were seeded in each well of 96-well plate and topped by dTHP1 suspension to obtain the ratio of interest. Spheroid culture in the presence of dTHP1 conditioned medium consisted of 5000 MDA-MB-231 cells in 10*µ*L of maintenance medium completed with 90*µ*L of dTHP1 supernatant.

Spheroid formation was monitored thrice a week under a bright field microscope, an AxioObserver (Zeiss) equipped with a Hamamatsu camera system. Images were acquired at 5× magnification. Only the wells with a single spheroid were used in our experiment.

Spheroid growth rates were estimated using the AreaShape_Area field (spheroid area in pixels) over the time of CellProfiler’s object identifier. The growth rate was estimated by linear regression of area vs time for each spheroid from 3 experiments.

### Gene expression profiling of spheroids

176 differentiated THP1 were isolated from co-cultures by FACS, processed using the Smartseq3xpress protocol ^123^. In addition, mono-cultures of differentiated or undifferentiated THP1 cells were processed as minibulks using the SmartSeq2 protocol ^124^.

To do so, 10,000 to 20,000 cells were pelleted by centrifugation and stored at −80°C. RNA were extracted using RNAeasy micro kit (Qiagen) following manufacturer instructions. The RNA were quantified using Qubit RNA high sensitivity (ThermoFisherScientific) and the integrity with a Bioanalyzer 2100 (Agilent). Samples with RIN *<* 8 were used for RNA sequencing using a modified SmartSeq2 protocol. Ten nanogram of RNA were denaturated in presence of 0.5uM of Oligo-dT30VN: (5’-AAGCAGTGGTATCAACGCAGAGTACT30VN-3’)and 0.5mM of dNTP for 10 minutes at 72 degrees. The reverse transcription used 25mM TrisHCl pH 8.3, 30mM NaCl, 0.5mM GTP, 2.5mM MgCl2, 8mM DTT, 0.5u/uL RNAse Inhibitor (Takara) 1uM TSO: (5’-AAGCAGTGGTATCAACGCAGAGTACATrGrG+G-3’). The reaction takes place in a thermocycler: 90minutes at 42 degrees, 10 cycles of 2 minutes at 50 degrees and 2 minutes at 42 degrees, ended by 5 minutes at 80 degrees. A second reaction using ISPCR oligo: (5’-AAGCAGTGGTATCAACGCAGAGT-3’) was conducted using 1X KAPA HiFi HotStart ReadyMix. The PCR amplification was: 3 minutes at 98 degrees, 10 cycles of 20 seconds at 98 degrees, 20 seconds at 67 degrees, 5 minutes at 72 degrees ended by 5 minutes at 72 degrees. The cDNA after dilution was tagmented for 10 minutes at 55 degrees using Amplicon Tagmentation mix (Tn5, Nextera). The reaction was stopped by the addition of 0.2% SDS. The product of this reaction was then indexed using Phusion HF as extensively described in the SmartSeq3 protocol.

Following deep sequencing, zUMIs version 2.9.7 was used to process raw FASTQ files. Reads were mapped to the human genome (hg38) using STAR version 2.7.3. The down-stream analyses were performed with R/Rstudio. Genes found in less than 3 cells were discarded. Differential expression analysis was performed using DESeq2 (test: MAST, significance threshold: 0.01).

FASTQ files are available on ArrayExpress (accession number: E-MTAB-12325).

### Immunostaining of mono-culture and co-culture spheroids

Spheroids were washed twice in phosphate buffer saline (PBS) and fixed with 4% Paraformaldehyde (PFA). The fixation was followed by a quenching step with 0.2M Glycine in PBS and two washes with 0.2% PBSTween20. A blocking step with 5% goat serum, 0.2% PBSTween20, 0.1M Glycine was performed prior to overnight incubation with primary antibodies in the same buffer. The next day the cells were washed twice with 0.2% PBSTween20 and stained overnight with the secondary antibodies and Hoechst33342 in the blocking solution. The next day the stained cells were washed twice in 0.2% PBSTween20 and once with PBS. Spheroids were cleared for 2 days in Omnipaque. The cleared structures were then transferred into silicon gasket covered with glass coverslips prior to imaging with a LSM980-AiryScan2 (Zeiss). Confocal images were acquired every 5*µ*m (Settings, 1AU for each fluorochrome) through the spheroids. All the images were converted from .czi to 8bit .tiff using ZenBlue software. The exported images were analyzed using CellProfiler4 and the results further analyzed in R/RStudio.

### Comparing cellular composition and gene expression in mouse vs human breast tumors

The cellular composition (fractional abundance) was log-transformed to avoid over-weighting highly abundant cell types in the PCA.

Because mouse and humans have distinct compositions of immune cells in general — more neutrophils in human vs more lymphocytes in mouse — the composition of mouse and human tumors was centered separately. Doing so allowed us to ask whether the variation in the immune composition of human and mouse tumors overlapped after accounting for sample-independent cross-species differences in immune populations. Principle components were computed by concatenating the mouse and human samples. Each sample was weighted as the inverse of the number of samples from that species (1/8 for mouse, 1/26 for human) so as to give equal weight to both species when computing principal components despite the difference in the number of samples in each species (n=8 for mouse, n=26 for human).

We further examined the gene expression dynamics of co-inhibitor and suppressive interleukin genes previously shown to associate positively with the progression of human lung tumors ^32^ (see Figure 3 of that article). To facilitate comparison to the bulk gene expression profiling of Mascaux et al. ^32^, we performed a pseudo-bulk by summing up, for each gene, the UMIs from the single cells of each gene and normalizing the fraction of UMIS to each gene of interest to a library size of one million (TPMs).

### Validating cell type abundance using HIFI

We used the spatial proteomic workflow HIFI by Watson et al. ^49^ to confirm the TME composition at the protein level.

HIFI was performed following the protocol described by Watson et al. ^49^. Briefly, slides with tissue sections were removed from the −80°C freezer, thawed at room temperature (RT) for 30 min, and washed in PBS for 1x 5 min. The sections were fixed with 4% PFA (Electron Microscopy Sciences) for 10 min on ice and washed 2x 5 min in PBS. Quenching was performed by incubating the sections with 0.1 M Glycine (Sigma-Aldrich) for 20 min at RT, followed by 2x 5 min wash in PBS. A hydrophobic border was drawn around the tissue section with a PAP pen. tissue permeabilization was performed with 0.2% Triton X-100 for 10 min at RT, followed by 3x 4 min PBS wash. Blocking (of non-specific antibody binding) was done by incubating the tissue sections with blocking buffer consisting of 10% normal donkey serum (Merck), 100 mM Ammonium chloride (NH4Cl) (Sigma-Aldrich), 150 mM Maleimide (Sigma-Aldrich) in PBS, for 1h at RT, in a humidified chamber. Blocking buffer was aspirated, and immediately replaced with the primary antibody dilutions in HIFI staining buffer consisting of 5% normal donkey serum, 4 mM NH4Cl in PBS, and incubated for 2.5h at RT. Tissue sections were washed 3x 5 min in PBS, and incubated with a secondary antibody solution in the HIFI staining buffer for 1h at RT. Primary antibodies were diluted at different concentrations (as shown in Table S9). All secondary antibodies were diluted 1:300, and DAPI was diluted 1:2000. The Secondary antibody mix was centrifuged at 10,000 RPM for 10 min before use. After secondary antibody incubation, sections were washed in PBS 3x 5 min at RT, excess PBS was removed, and slides were mounted with SlowFade Diamond Antifade Mounting Medium (ThermoFisher S36963).

Imaging was started immediately after mounting, using Axio Scan Z1 scanner. In the first imaging round, tissue detection was done in order to identify regions of interest (ROIs) which were reused for each subsequent cycle.

The next day, after imaging was complete, coverslips were removed by gently rocking the slides in PBS at RT. Tissue sections were subsequently wash 3x 5 min, and incubated with Elution buffer, consisting of 0.5 M Glycine (Sigma-Aldrich), 3 M Guanidinium chloride (Sigma-Aldrich), 3 M Urea (Sigma-Aldrich) and 40 mM Tris(2-carboxyethyl) phosphine hydrochloride (TCEP) (Sigma-Aldrich) in ddH20, for 3 min at RT. Samples were washed 3x 5 min at RT, and a new cycle of staining would be initiated, starting with the blocking step.

Washing, antibody incubations, and elution steps were performed on a 3-axis orbiter (with a shallow tilt angle).

This protocol was repeated as many times as it was needed in order to stain for all the protein targets of interest, as shown in Table S9. The last round of staining was performed only with DAPI and was used for background subtraction as detailed below.

For the earliest time point (day 11 post-tumor injection), as tumors were too small to be used for both sequencing and imaging, tumors’ fixed and frozen tissue sections of different mice from the same cohort were used. For other time points (day 14, 18, and 24 post-tumor injections), as the tumor size was sufficient, scRNAseq and imaging were performed on tumors originating from the same mouse. The following cell type markers were used to perform the cell type calling: SOX9 (cancer cells), CD45 (immune cells), CD3-AF750 (T cells), CD8a-647 (cytotoxic T cell), Iba1 (myeloid cells), F4/80/Adgre1 (macrophages), CD68 (phagocytic macrophages), and DAPI to stain cell nuclei.

The images obtained using the Zeiss Axio Scan Z1 microscope were post-processed using the Zeiss Zen Blue software platform to perform tile stitching and fusion on all images. To remove the background fluorescence, the rolling-ball method implemented in the Zen software (using a diameter of 75 µm) was applied. As the sample is stained in several rounds, images from each staining round need to be aligned/registered together to obtain the final multiplex image: the registration of stitched images was performed using the imreg dft package, an implementation of discrete Fourier transformation-based image registration ^125^. Once the images were registered, a manual verification was conducted to identify possible mis-stitched areas. These areas were manually identified and annotated using QuPath v0.3.2 ^126^ to be ignored in the downstream analysis. Some areas of the tissue can also get damaged during the staining and imaging. To avoid analyzing these areas, all cells missing one round of DAPI staining were automatically annotated and excluded from downstream analysis using an in-house Python script.

The resulting OME TIFF images – containing all the staining rounds – still featured some nonspecific background signals, such as strong fluorescence on the tumor border for certain channels. To eliminate this artifact that could bias cell type identification, we subtracted the last round of imaging (round 7, times a constant) from all the preceding imaging rounds: the last round of imaging was a control round in which no primary antibody was used. Once the images’ sufficient quality was confirmed, cells were segmented using a manually pre-trained CellPose model ^127^.

Following cell segmentation, we determined the type of each cell. One challenge in doing so is that different cell type markers have different levels of brightness and dynamic range. Simply clustering cells into cell types according to marker expression leads to clustering cells according to most variable markers, which masks cell types detectable by less variable markers. To address this, markers’ expression needs to be scaled before clustering. One challenge with scaling this type of data lies in the bimodal nature of the log-transformed fluorescence intensities. By nature, the fluorescence intensity distributions are bimodal and feature different proportions of positive to negative cells for each marker. For example, about half of the cells express the cancer cell marker, but very few cells express the cytotoxic T marker, especially at an early time point. In the case of the rare cell population, simply centering and scaling – dividing by the standard deviation – centers the distribution on the negative cell population, as they represent the vast majority of cells, and amplifies the signal of cells weakly expressing the marker, as if these cells were positive for it. To avoid this bias, we choose to scale the marker’s intensity by using the highest mode of the multimodal distribution of each marker across samples. Rare populations required different smoothing parameters to identify the different Gaussians composing the Gaussian mixture compared to frequent cell populations, as the positive cells are very few, making it difficult to distinguish the Gaussian mixture responsible for positive cells from the noise. To overcome this difficulty, we proceeded by subpopulation: we looked for the CD3 or CD8 high mode only among CD45+ cells for example.

Scaling markers in this way results in the negative cells having values close to 0 while the positive cell population log-transformed fluorescence values centered around 1.

After markers were scaled, cells expressing at least one marker were clustered using Leiden clustering on the two first PCs of the log-transformed scaled fluorescence signals (number of neighbors *k* = *max*(*round*(*nb_cells_* × 10*^−^*^3^), 10), and resolution of *res* = 1) to identify the cell composition of each sample. Performing PCA helped cluster cells into cell types consistently across images using the same parameters *k* and *res* above. PC1 distinguishes between cancer versus immune cells and PC2 distinguishes immune cell types (myeloid versus lymphoid). Additional PCs explained very little of the variance in marker abundance and appeared to confuse the Leiden clustering more than helping it to resolve cell types (Figure S10). Each cluster was then automatically annotated to one cell type using a list of necessary markers and possibly expressed markers (Table S10). Finally, log-transformed cell type proportions obtained from HIFI data were compared to the ones obtained from the single cell analysis for day 14, 18, and 24 samples.

## Acknowledgements

The authors thank members of the Joyce and Hausser labs for critical discussion, and specifically Jakob Rosenbauer, Petter Säterskog, Anissa El Marrahi, Andreas Lundqvist, and Vladimir Wischnewski for their regular input on the project.

We thank Bastien Mangeat, Elisa Cora, Lionel Ponsonnet, and Marion Leleu from the EPFL Gene Expression Core facility for input on the experimental design, for performing the single-cell gene expression experiment, and for help with the initial gene expression analysis. The authors acknowledge the support from the Swedish Cancer Society, the Swedish Research Council, SciLifeLab, and Karolinska Institutet (all to J.H), and the Cancer Research UK Grand Challenge Grant, the Breast Cancer Research Foundation, Ludwig Institute for Cancer Research, and University of Lausanne (all to J.A.J.). S.S.W. was supported by The Brain Tumor Charity Future Leaders Fellowship, and M.M. was supported by funding from the AIRC and European Union’s Horizon 2020 research and innovation program under the Marie Sklodowska Curie (grant agreement 800924).

## Author contributions

S.S.W., J.A.J. and J.H. conceived the study and designed experiments. J.H. and I.S. performed the experiments with support from S.S.W., M.M., K.S. and J.A.J.. L.G. and J.H. analyzed and interpreted the single-cell RNAseq data with input from S.S.W., K.S., A.C., J.E.M, and J.A.J.. L.G. and J.H. wrote the manuscript, and S.S.W and J.A.J. edited the manuscript. J.A.J. and J.H. acquired funding. All authors reviewed or commented on the manuscript.

## Data availability

Source data files to reproduce the analyses can be downloaded at GEO (accession GSE221528): https://www.ncbi.nlm.nih.gov/geo/query/acc.cgi?acc=GSE221528, reviewer token ‘khmxiwsmrbqdbux’.

## Code availability

The code to reproduce the analyses can be downloaded at https://data.mendeley.com/preview/82k8g6y3hf?a=16770d6a-f4e3-4114-9098-fed42f80b218.

