## Supplemental data for "Multi-cellular phenotypic dynamics during the progression of breast tumors"

### Supplemental Tables

#### Supplementary Table 1

Table S1: Comparing average transcriptional profiles of the 9 single-cell RNA sequencing (scRNAseq) clusters in our experiment with bulk RNAseq profiles from FACS-isolated cell populations confirms the cell types identified by marker gene analysis. For each scRNAseq cluster, we list the five FACS-ed populations with the highest correlation. NIH_3T3 cells are mouse embryonic fibroblasts; C1C12 is an immortalized mouse myoblast cell line; C3H_10T1_2 is a murine embryo fibroblast cell line; 3T3-L1 cells are preadipocytes; MEFs are fibroblasts; RAW_264_7 is a monocyte/macrophage cell line.

| cluster | top 5 correlations |
|---|---|
| cancer cells | nih_3T3 (r=0.67) ; C2C12 (r=0.67) ; C3H_10T1_2 (r=0.65) ; 3T3-L1 (r=0.65) ; MEF (r=0.64) |
| myeloids | macrophage_bone_marrow_2hr_LPS (r=0.67) ; macrophage_bone_marrow_0hr (r=0.66) ; osteoclasts (r=0.66) ; microglia (r=0.65) ; macrophage_bone_marrow_6hr_LPS (r=0.65) |
| DCs | macrophage_bone_marrow_2hr_LPS (r=0.64) ; macrophage_bone_marrow_0hr (r=0.64) ; osteoclasts (r=0.63) ; RAW_264_7 (r=0.63) ; dendritic_cells_myeloid_CD8a- (r=0.62) |
| macrophages / monocytes | macrophage_bone_marrow_2hr_LPS (r=0.67) ; macrophage_bone_marrow_0hr (r=0.66) ; osteoclasts (r=0.66) ; microglia (r=0.66) ; macrophage_bone_marrow_6hr_LPS (r=0.65) |
| lymphoids | CD4.24h.LN#1(r=0.62) ; nih_3T3 (r=0.62) ; macrophage_bone_marrow_2hr_LPS (r=0.62) ; macrophage_bone_marrow_0hr (r=0.62) ; CD8.24h.LN#1 (r=0.62) |
| conventional T cells | CD4.24h.LN#1 (r=0.63) ; nih_3T (r=0.62) ; CD8.24h.LN#1(r=0.62) ; RAW_264_7 (r=0.62) ; macrophage_bone_marrow_2hr_LPS (r=0.62) |
| Th cells | CD4.24h.LN#1(r=0.63) ; nih_3T3 (r=0.62) ; CD8.24h.LN#1(r=0.62) ; RAW_264_7 (r=0.62) ; CD4.24h.LN#2 (r=0.62) |
| Tc cells | CD4.24h.LN#1 (r=0.63) ; nih_3T3 (r=0.63) ; CD8.24h.LN#1 (r=0.62) ; RAW_264_7 (r=0.62) ; macrophage_bone_marrow_0hr (r=0.62) |
| Tregs | CD4.24h.LN#1 (r=0.63) ; CD8.24h.LN#1 (r=0.62) ; macrophage_bone_marrow_2hr_LPS (r=0.62) ; nih_3T3 (r=0.62) ; macrophage_bone_marrow_0hr (r=0.62) |
| group1 ILC | nih_3T3 (r=0.63) ; NK.CD49b+.Lv#3 (r=0.62) ; RAW_264_7 (r=0.62) ; macrophage_bone_marrow_0hr(r=0.62) ; macrophage_bone_marrow_2hr_LPS (r=0.62) |
| B cells | nih_3T3 (r=0.63) ; macrophage_bone_marrow_2hr_LPS (r=0.63) ; macrophage_bone_marrow_0hr (r=0.63) ; RAW_264_7 (r=0.63) ; C2C12 (r=0.62) |
| stromal cells | osteoblast_day5 (r=0.64) ; osteoblast_day14 (r=0.64) ; 3T3-L1 (r=0.64) ; MEF (r=0.63) ; osteoblast_day21 (r=0.63) |

#### Supplemental Table 2

**Table 2** — Gene and pathway ParTI enrichment analysis on cancer cells

#### Supplemental Table 3

**Table 3** — Gene and pathway ParTI enrichment analysis on monocytes & macrophages

#### Supplemental Table 4

**Table 4** — DESeq2 analysis of 3D co-culture of human breast cancer cells and macrophages vs 3D mo-culture of macrophages

#### Supplemental Table 5

**Table 5** — FGSEA analysis of PC loadings for Tc cells

#### Supplemental Table 6

**Table 6** — FGSEA analysis of PC loadings for ILCs

#### Supplemental Table 7

**Table 7** — FGSEA analysis of PC loadings for Th cells

#### Supplemental Table 8

**Table 8** — FGSEA analysis of PC loadings for Tregs

#### Supplemental Table 9

**Table 9** — HIFI markers' detailed information

#### Supplemental Table 10

**Table 10** — Cell type x markers for HIFI cell type automated annotation

#### Supplemental Figures

### Supplementary Figure 1

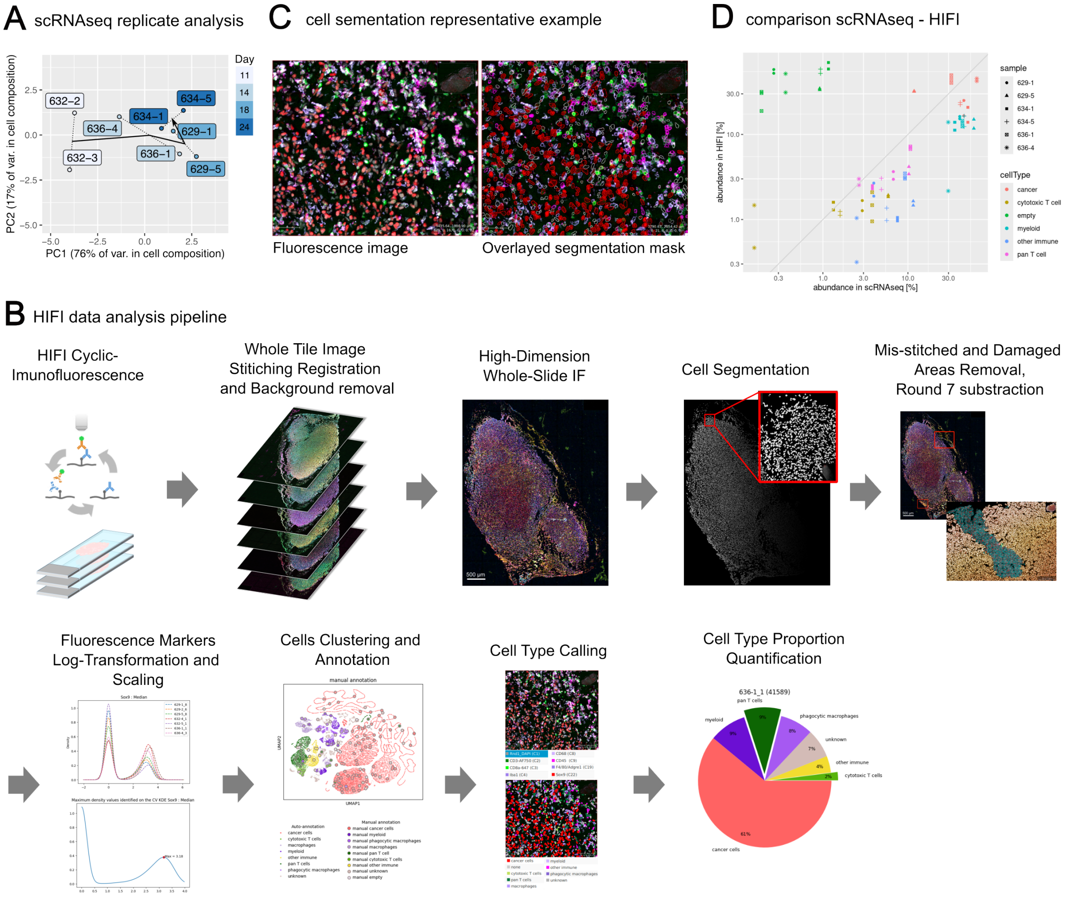

Figure S1: **A**. Tumors from different animals follow a similar temporal path in terms of cellular composition dynamics. We performed principal components analysis (centered, unscaled) on the log fractional abundance of cell types from scRNAseq samples from different time points and biological replicates. **B**. Overview of the hyperplexed immunofluorescence imaging (HIFI) workflow. Staining and imaging were followed by image stitching, registration, and background removal to create multi-dimensional images. Cells were segmented using CellPose with a manually pre-trained model. Mis-stitched regions were removed by visual inspection. Cells located in tissue regions damaged or lost at some point during the staining and imaging rounds were excluded by setting a minimal threshold on DAPI across imaging rounds: if a cell lacked DAPI in at least one imaging round, it was deemed located in a lost tissue region. The fluorescence intensity signal was log-transformed and scaled so that a value of 0 corresponded to negative staining and a value of 1 to positive staining (Methods), in order to facilitate unsupervised clustering. Cells were then clustered using Leiden on the two first principal components (PCs) of log-transformed scaled fluorescence markers. Each cluster was automatically annotated as an individual cell type by matching to a dictionary of cell-type-specific markers (Methods). Cell type proportion was then computed for each slide of each sample. **C**. The representative outcome of cell segmentation by CellPose on sample 636-1 section 1, day 18 post tumor injection. **D**. The abundance of cell types measured by scRNAseq matches the abundance of cell types determined by HIFI. One cell type (empty) had much stronger abundance in HIFI. This is due the absence of positive markers for a large proportion of cells of the HIFI data. In contrast, genome-wide gene expression profiling by scRNAseq allowed assigning a cell type to the vast majority of cells. Because cell typing could be done reliably for more cells in scRNAseq, the abundance of cell types appears higher in scRNAseq compared to HIFI. Beyond these methodological differences, cell types highly abundant in scRNAseq were also highly abundant in HIFI - cancer, myeloid. Similarly, cell types of lower abundance in scRNAseq were also found at lower abundance in HIFI - cytotoxic T cell, pan T cell, and other immune.

### Supplementary Figure 2

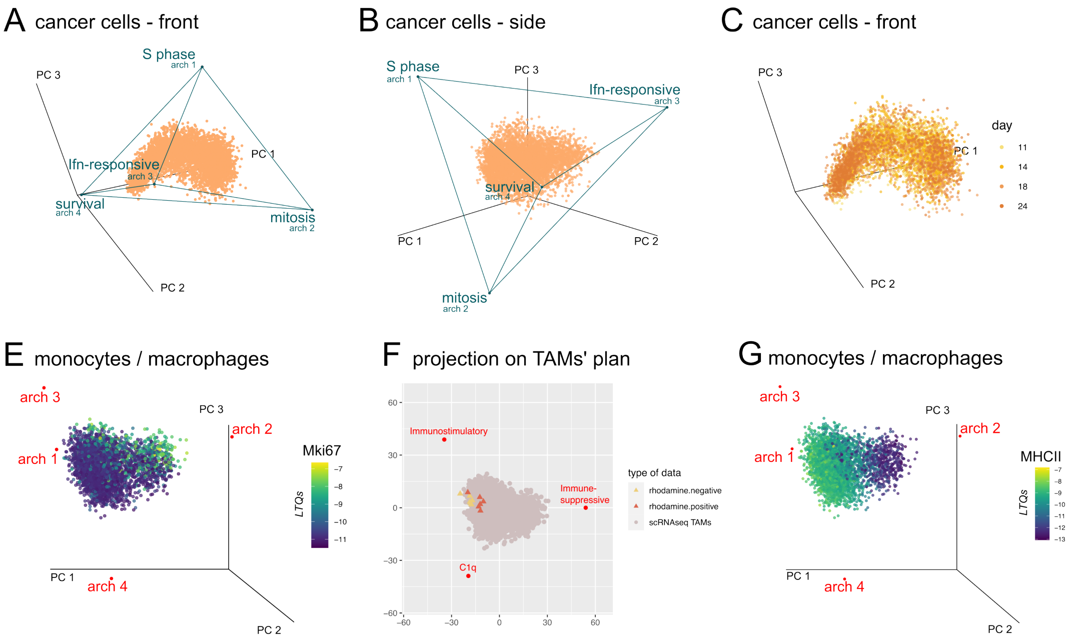


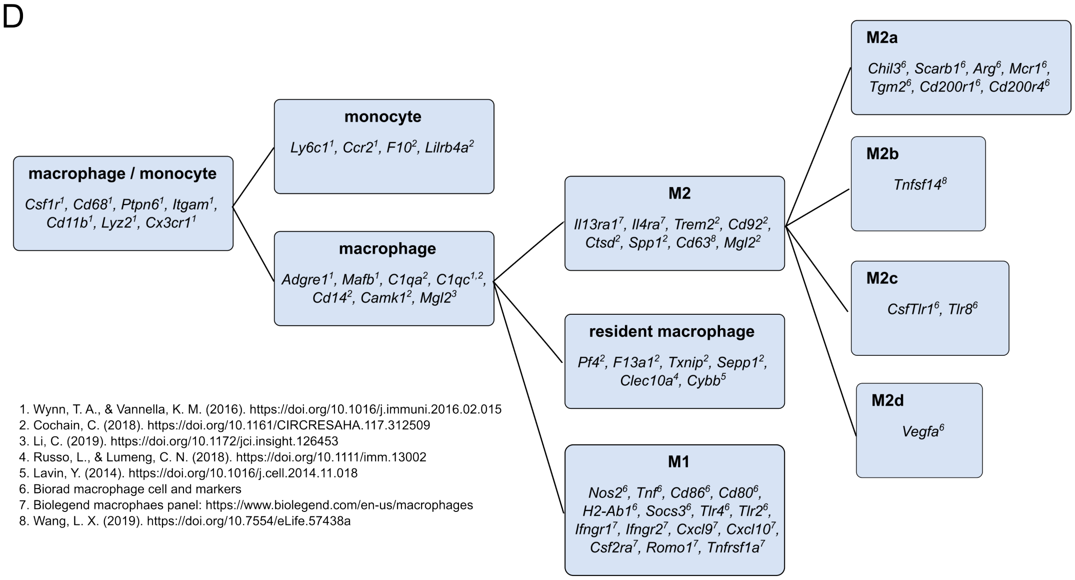


1. Wynn, T. A., & Vannella, K. M. (2016). https://doi.org/10.1016/j.immuni.2016.02.015
2. Cochain, C. (2018). https://doi.org/10.1161/CIRCRESAHA.117.312509
3. Li, C. (2019). https://doi.org/10.1172/jci.insight.126453
4. Russo, L., & Lumeng, C. N. (2018). https://doi.org/10.1111/imm.13002
5. Lavin, Y. (2014). https://doi.org/10.1016/j.cell.2014.11.018
6. Biorad macrophage cell and markers
7. Biolegend macrophaes panel: https://www.biolegend.com/en-us/macrophages
8. Wang, L. X. (2019). https://doi.org/10.7554/eLife.57438a

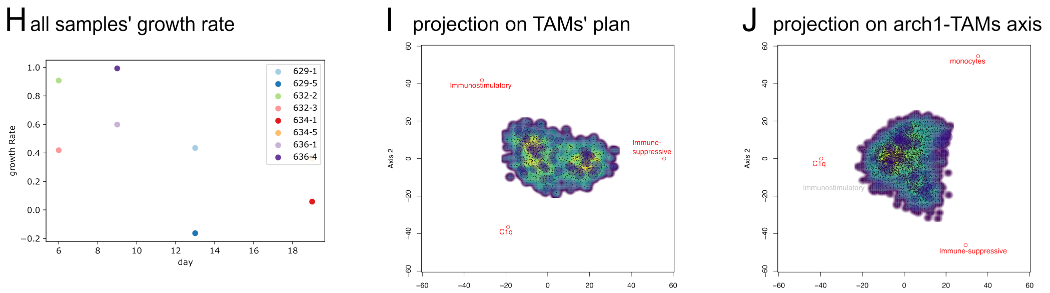

Figure S2: **A-C**. Fitting a tetrahedron on cancer cells cluster using the ParTI method[1] provides a framework for gene and pathway enrichment analysis (see Table S2). Dots represent cancer cells in PCs space. Axes represent the three first PCs. The same projection is used in panels Figure 2 A-D. **C**. The proliferation – survival phenotypic continuum in cancer cells is consistently present across the time span of the experiment. Colors: days post tumor injection. **D-E**. The density of single tumor-associated macrophages (TAMs) in gene expression space is best described as a continuum. Ellipses represent the error in gene expression estimated by Sanity. Colors represent the density of cells (dark blue is low density, yellow is high density). Dots represent the 33% of monocytes/macrophages closest to the plane defined by the M1-like/immunostimulatory, C1q and M2-like/immune-suppressive archetypes (panel **D**) or all cells projected onto the monocyte-C1q-immune-suppressive plane (panel **E**). **F**. Genes signatures for monocyte and macrophage phenotypes were manually chosen by reviewing the literature. **G-H**. Cells close to archetype 2 (Arch2) – immune-suppressive phenotype – upregulate proliferation markers and down-regulate (major histocompatibility complex class II (MHCII))–dependent antigen presentation. Dots: single macrophages and monocytes. Axes: three first principal components. Color bar: gene expression (log transcription quotients (LTQs)). **I**. Perivascular macrophages associate more with the immune-suppressive archetype than non-perivascular macrophages. **J**. Sample 629-5 from time point 14 is a tumor that spontaneously receded, unlike other samples.

### Supplementary Figure 3

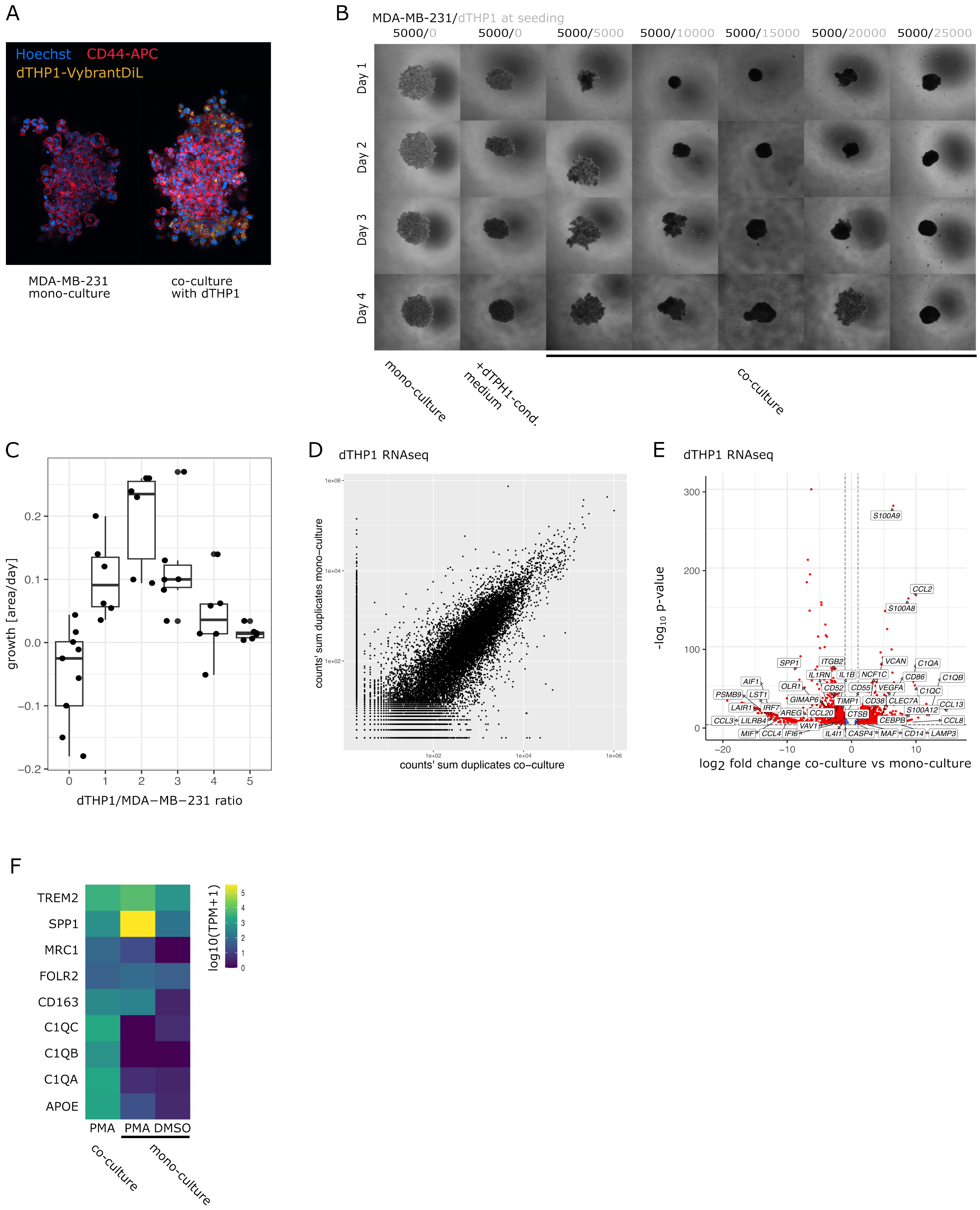

Figure S3: 3D co-culture of human breast cancer cells and macrophages validate cancer-macrophage cross-talks suggested by single-cell gene expression profiling of progressing tumors in mice. **A**. 3D co-culturing MDA-MB-231 human breast cancer cells with the THP1 human macrophage cell line differentiated by PMA leads to infiltration of tumor spheroids by macrophages. CD44 is a cell adhesion marker. **B**. Co-culturing breast cancer cells with differentiated macrophages induces changes in spheroid growth and compactness. Bright-field imaging of spheroids seeded with different numbers of cancer cells and macrophages (top labels). Change in compaction is due both to macrophages and diffusible signals: adding culture medium from dTHP1 cells to cancer mono-cultures increased the compactness of the spheroid but to a lesser extent than co-culture. **C**. Spheroid growth depends on the amount of macrophages in the co-culture. Each dot represents a spheroid. The growth rate was estimated by quantifying temporal changes in the spheroid area (pixels) over time by longitudinal microscopy and image analysis. **D**. Correlation between 'gene log10 expression (sum by genes of the replicates counts) in a 3D mono-culture of dTHP1 cells and a 3D co-culture of dTPH1 cells with cancer cells. The overall expression correlates, but some genes have a clear differential expression (as highlighted in panel D). **E**. Differential gene expression analysis upon exposing dTHP1 cells to cancer cells shows up-regulation of C1QA, C1QB, and C1QC. **F**. Archetype 4-defining genes C1Q and APOE are induced by exposing THP1 to cancer cells, not upon differentiation by PMA alone.

### Supplementary Figure 4

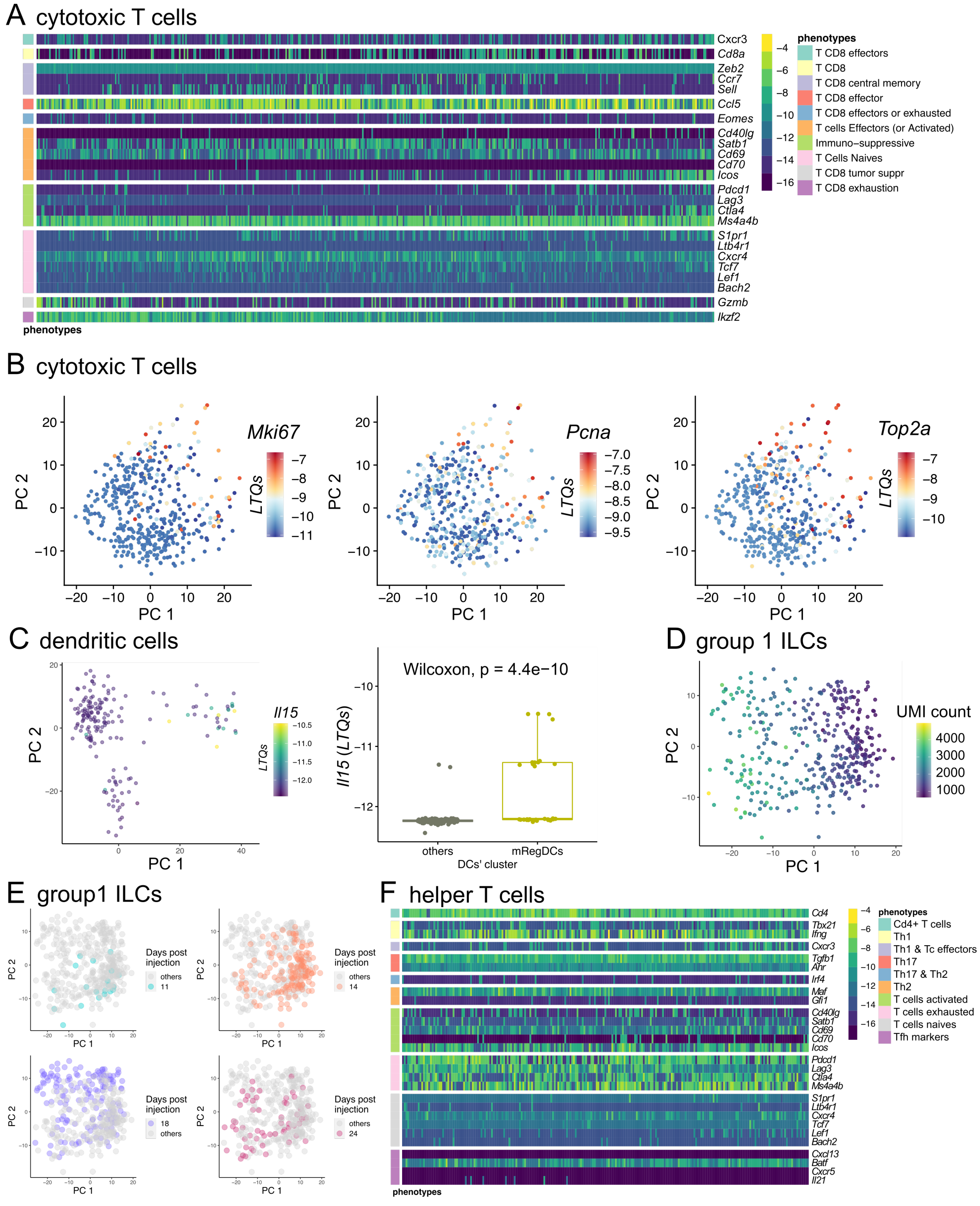

Figure S4: **A**. State-of-the-art gene signatures used to describe cytotoxic T cells (Tc) phenotypic heterogeneity poorly fit the transcriptional heterogeneity in the present experiment. For example, the naive and memory T cell marker *Cd7/Gp40* [2] is expressed in the Xcl1+ cluster, which also expresses activation markers *Satb1* and *Ikzf2/Helios* [3] (Figure S4A). *Satb1*, *Ikzf2/Helios* and *Icos* are all markers of effector T cells, but they are expressed by different clusters here (Figure S5). To further highlight the mixed naive/memory/effector phenotypes of these cells, naive T cell markers such as *S1pr1*, *Tcf7* and *Cxcr4* are detectable in all the cells, and so are the effector markers *Cd69* and *Ccl5*. As for the central memory markers *Zeb2*, *Ccr7* and *Sell*, we find expressions in mutually exclusive sets of cells. Columns: Tc cells. Rows: conventional Tc phenotypic markers. Color bar: gene expression (LTQs). **B**. Tc phenotypic heterogeneity is partially explained by a gradient of proliferation that follows the cytotoxicity/activation gradient (Figure3 D). **C**. mregDCs express *Il15* highest compared to classical dendritic cells (labeled "others" on the figure). **D**. Group 1 innate lymphoid cells (ILCs) (dots) that score low on PC1 (identified as innate lymphoid cell type 1s (ILC1s) Figure3 J) have a high number of unique molecular identifier (UMI) per cell, indicating a high transcriptional activity. **E**. Group 1 ILCs temporal phenotypic dynamics mainly follows the gradient of activation captured by PC2 (Figure3 J). **F**. While the phenotypic heterogeneity of T helper cells (Th) is hardly explained by known phenotypic marker genes, a Th1 signature fits Th cells best. The Th1-marker *Ifng* is highly expressed (LTQs $> -8$, lowly expressed genes have LTQs $< -10$, genes with highest expression have LTQs $> -6$) in 40% of Th cells (Figure S4C). The Th1 transcription factor *T-bet* is detectable in many cells (Figure S4C), as well as Th1 effector genes (see results in the main text). Cell types associated with Th1 response such as Natural Killer (NK) and ILC1s (see results) are found in our samples. While Th cells express the T follicular helper (Tfh) transcriptional factor *Batf*, a Tfh phenotype does not fit here: neither the B-cell zone homing receptor *Cxcr5* nor the *Il21* B-cell stimulating cytokine nor the *Cxcl13* B-cell chemoattractant are detected.

#### Supplementary Figure 5

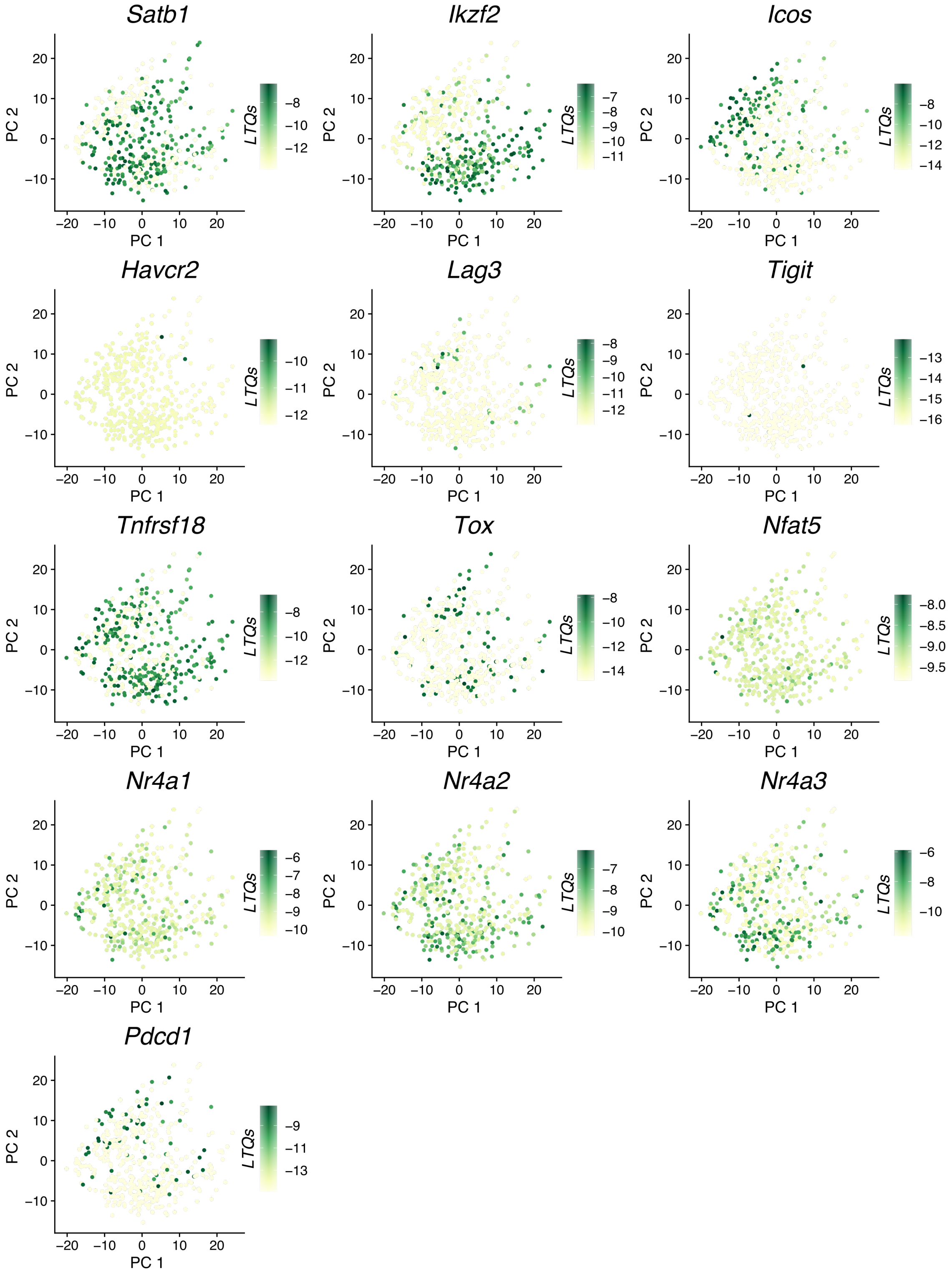

Figure S5: Tc phenotypic heterogeneity is not explained by an exhaustion gradient. Instead, exhaustion markers are expressed inconsistently across cells, except for *Icos* and *Ikzf2* whose expression is mutually exclusive. Dots: single Tc cells. Axes: first two principal components of Tc gene expression. Color bar: marker gene expression (LTQs). Genes are considered highly express when LTQs $> -6$ and not expressed when LTQs $< -12$.

### Supplementary Figure 6

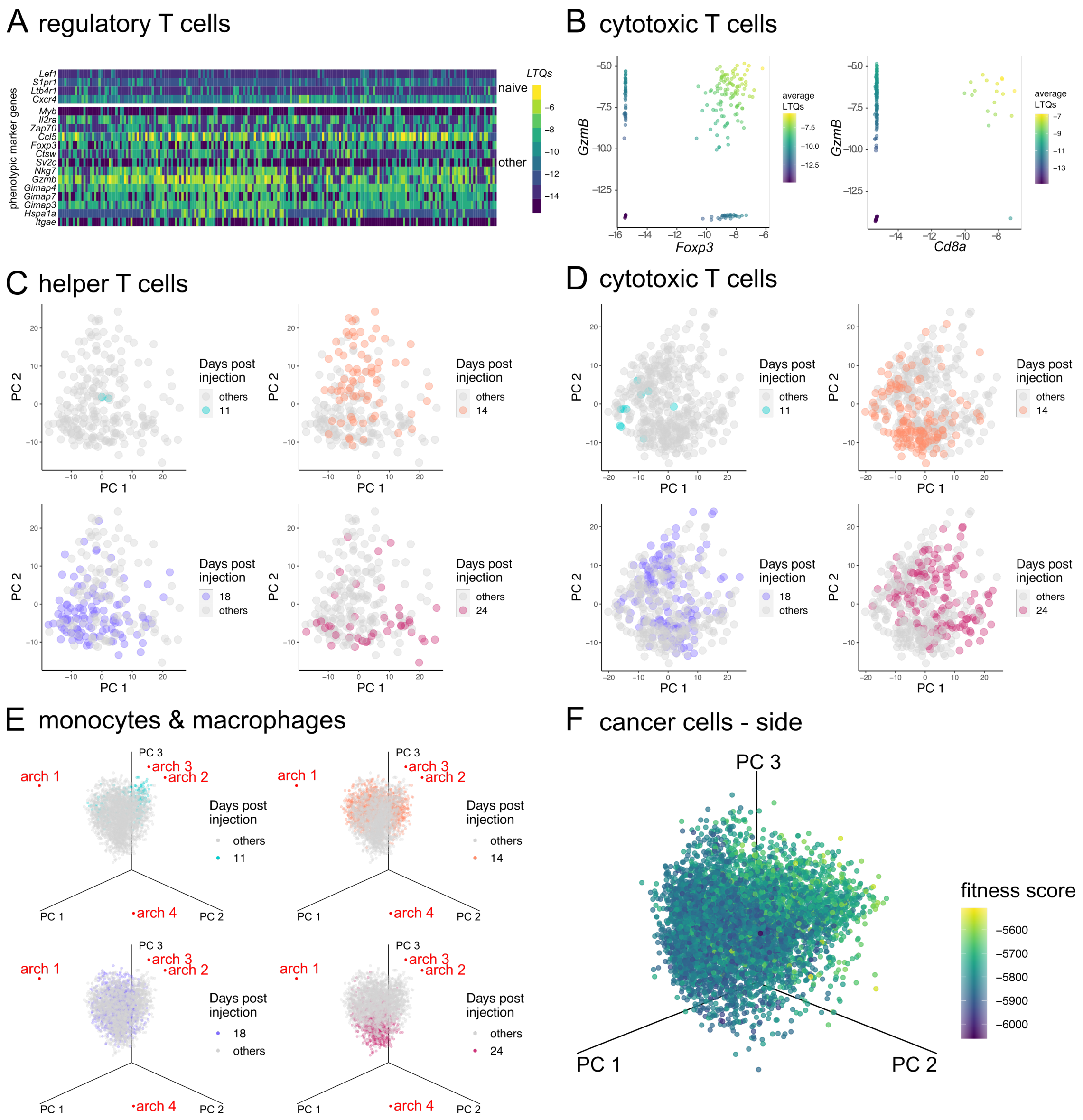

Figure S6: Cell phenotypic heterogeneity varies in association with the growing tumor. **A**. Regulatory T cells (Tregs) express several naive T cells markers and highly express *Gzmb*. **B**. Three observations exclude a mislabeling of Tc as Gzmb+ Tregs as (a) Tregs form a clearly distinct transcriptional cluster from *Cd8+* cells in our data (Figure1 B), (b) Tregs where *Cd8* is undetectable strongly express *Gzmb* and (c) *Gzmb* is highly expressed in *Foxp3*-expressing Tregs. Axes: gene expression (LTQs). Dots: single Tregs. **C**. Th phenotypic dynamics mainly take place along PC2, a matrisome / antigen presentation versus proliferation axis (Figure3 H). **D**. Tc score is initially low on PC1 and PC2 (day 11) and gradually shifts to a PC1/2 high phenotype, following a gradient of activation and cytotoxicity (Figure3 D). **E**. The prevalence of the C1q macrophage phenotype increases during tumor progression. **F**. Interferon (IFN)-responsive cancer cells have the highest fitness among cancer cells. Dots: single cancer cells from all time points. Color bar: cellular fitness score computed from the CRISPR screening data of Lawson et al.[4].

#### Supplementary Figure 7

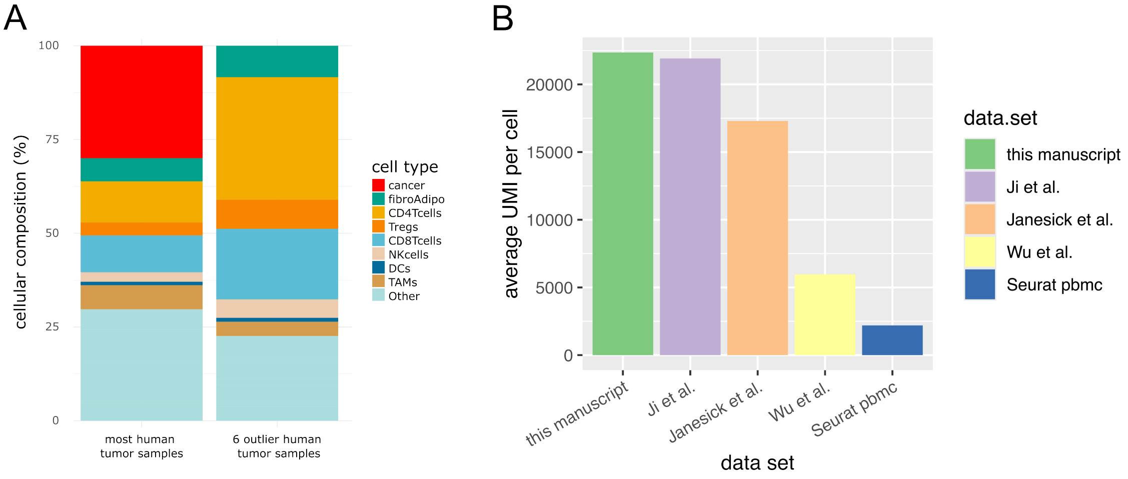


Figure S7: **A.** Outlier human breast tumors in terms of the principal components analysis of the cellular composition of (Figure 5A) are explained by an absence of cancer cells. Cellular composition of 20 human breast tumors and of the 6 outlier tumors. Cell types were obtained from the original study ( Wu et al. [5]). **B.** Sequencing depth of the scRNAseq data of this manuscript compared to state-of-the-art publicly available datasets.

### Supplementary Figure 8

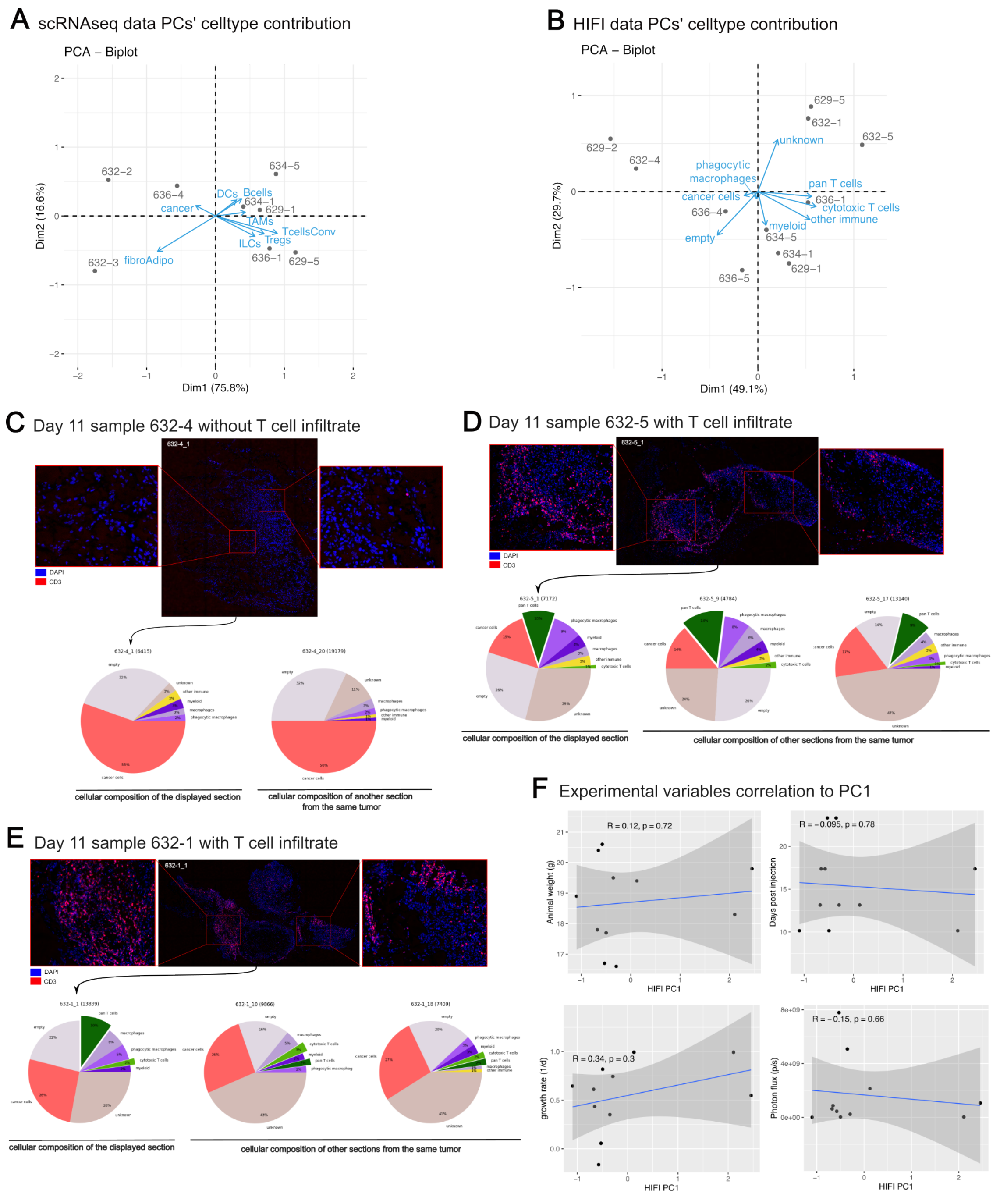

Figure S8: Across cellular abundance profiling techniques (scRNAseq vs HIFI), we find the same cancer – adaptive immunity dominant axis of variation across individuals. Yet, there is an important individuality in how rapidly tumors progress along this axis. **A.** Most of the inter-tumor variation in cellular abundance as profiled by scRNAseq is explained by PC1 (65.4%) and is driven by cancer – adaptive immunity axis. **B.**. The same axis of cancer – adaptive immunity is found as the main driver of cellular composition heterogeneity using HIFI data (protein level). **C-E.** Profiling tumor samples collected on the same day shows important individual differences in the dynamics of immune infiltration. **C.** Sample 632-4, collected at day 11 post tumor injection, is free from T cell infiltrate, consistent with observations from scRNAseq. **D-E.** Samples 632-5 and 632-1, also collected at day 11 post-tumor injection already feature important T cell infiltration typical of later scRNAseq time points. **F.** The origin of this inter-individual variation is still to be determined. Inter-individual variability in cellular composition as assayed by HIFI does not significantly associate with animal-specific variables such as tumor size (assayed by luciferase photon flux), tumor growth rate, animal weight, and time post-injection.

### Supplementary Figure 9

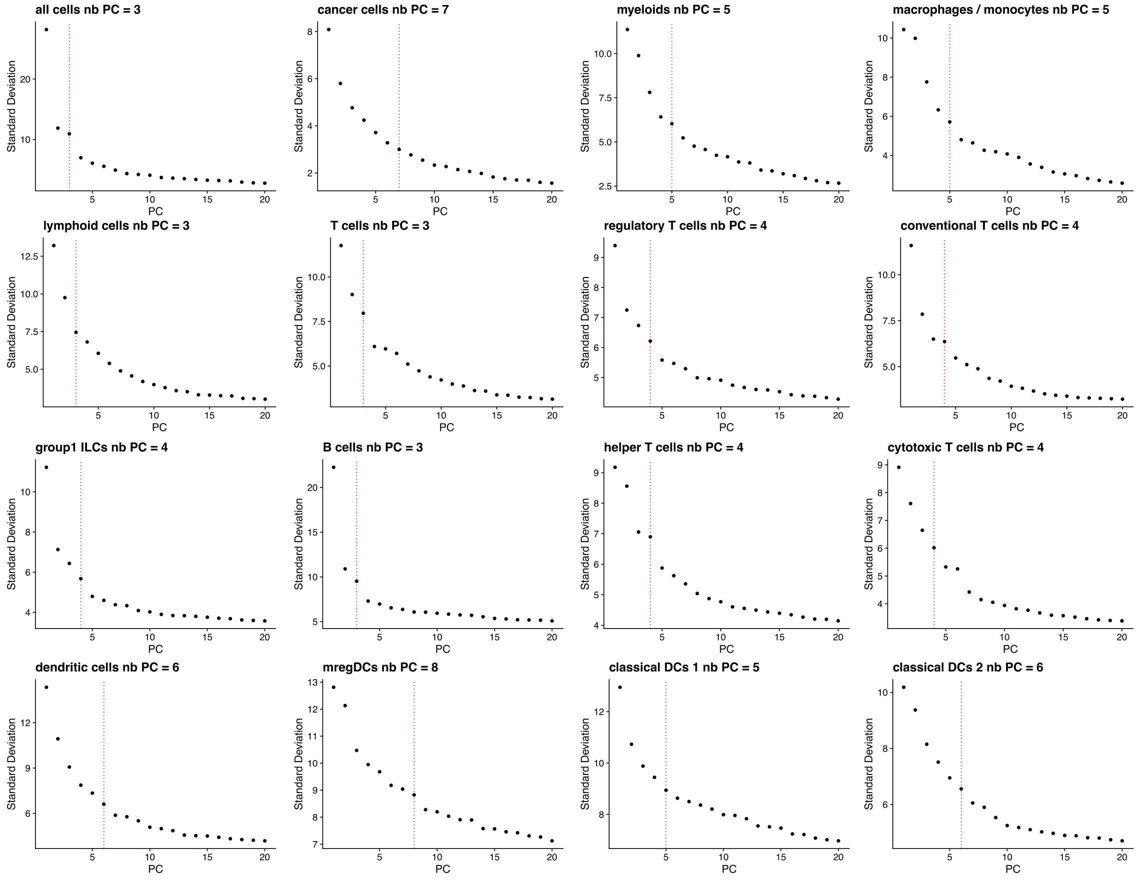


Figure S9: The number of PCs used to describe the transcriptional heterogeneity of different cell types was chosen using the elbow criteria. Inspecting the standard deviation in gene expression explained by each PC (y-axes) as a function of the number of PCs (x-axis), we chose a number of components beyond which adding further components provided comparatively little benefit in capturing transcriptional heterogeneity.

### Supplementary Figure 10

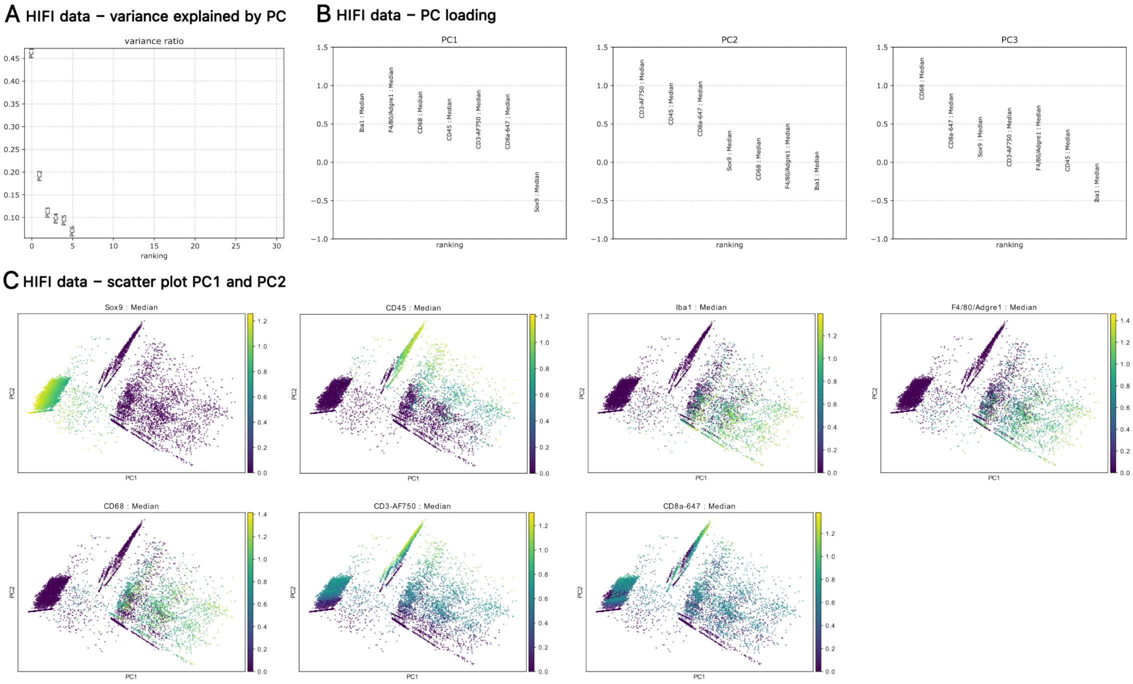


Figure S10: Performing principal components analysis (PCA) helped cluster cells into cell types consistently across images using the same parameters k and res above. **A.** The two first PCs explain more than half of the variance in the data. **B.** PC1 distinguishes between cancer and immune cells, when PC2 distinguishes immune cell types: myeloid and lymphoid. PC3 explained little of the variance in marker abundance and didn't help to distinguish cell types. **C.** The cell's projection in the first two PCs illustrate clearly how the main cell types (cancer, myeloid, lymphoid) separate in this space.

### Supplementary Figure 11

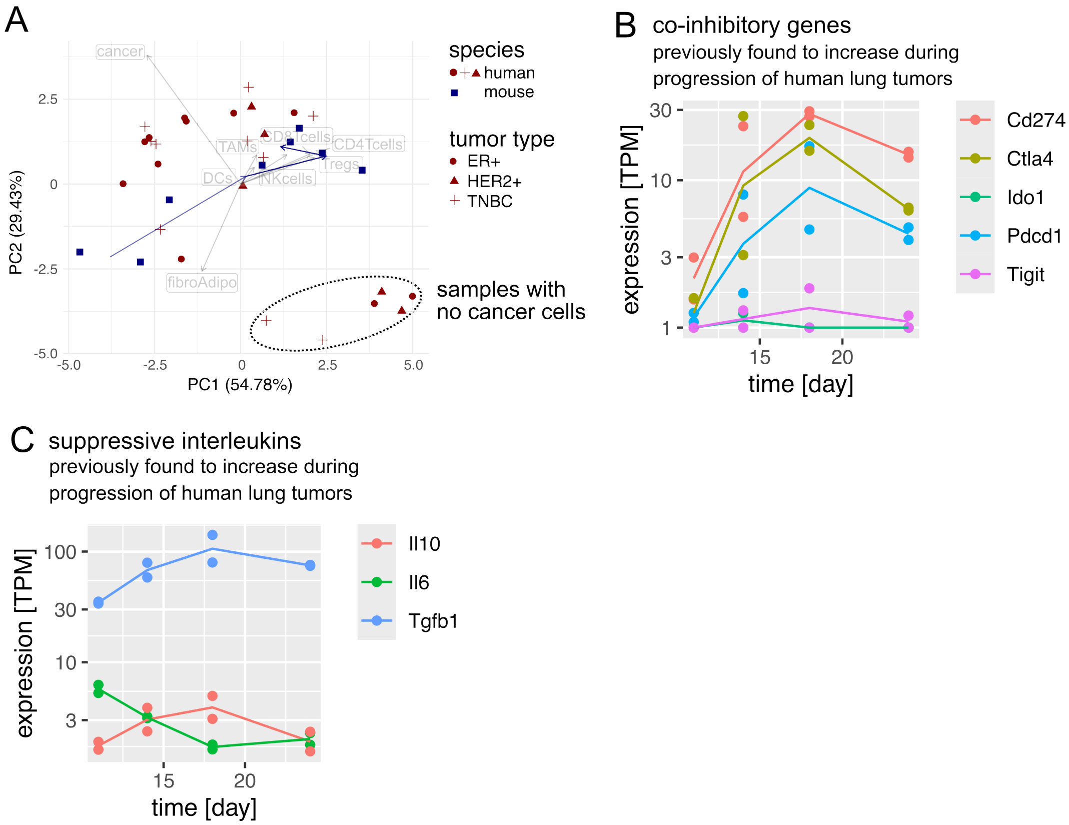


Figure S11: The cellular and gene dynamics of mouse tumors align with human breast tumors. **A.** Temporal variation in the cellular composition of progressing mouse breast tumors overlaps with inter-patient variation of human breast tumors. Principal components and bi-plot analysis of the cellular composition of 8 progressing mouse tumors and 26 human tumors (scRNAseq). Numbers: day (11, 14, 18, 24) of tumor collection post-induction. Blue arrow: average time trajectory of mouse tumors. Grey arrows: contribution of each cell type to the principal components. 6 human tumors appeared as outliers (circled), which finds an explanation in the absence of annotated cancer cells in these samples (Figure S7). **B-C.** Co-inhibitor (**B**) and suppressive interleukin (**C**) genes whose expression increases during the progression of human lung tumors[6] also increase in mouse breast tumors, albeit with a peak 18 days post tumor induction.

### Supplementary Notes

#### Supplementary Note 1: cellular and gene dynamics in mouse overlap with inter-sample variation of human breast tumors

To probe the relevance of cellular dynamics in our mouse model, we compared these dynamics to inter-tumor variation in humans.

Joint PCA of our mouse tumor scRNAseq with human breast tumor scRNAseq suggests that most inter-tumor variation in humans overlaps with the cellular dynamics of mouse tumors (Figure S11A). Both in mice and in humans, inter-tumor variation in cellular composition was driven by a cancer-adaptive immunity axis, with adaptive immune cells more present at later time points and cancer cells more prevalent in early mouse tumors. This trend was robust across human breast tumor subtypes. The cellular composition of early mouse tumors (day 11, Figure S11A) was not observed in human data, consistent with rare resection of early human breast tumors: 90% of human breast tumors are larger than 10mm[7] the largest size of our mouse tumors. A group of outlier human tumor samples did not align with mouse time dynamics due to an absence of cancer cells (Figure S7A).

Beyond cellular composition, gene expression patterns also have parallels between human and mouse tumors.

For example, Mascaux et al.[6] previously explored temporal gene regulation in progressing lung human tumors by ordering tumors by histological stage and performing bulk RNA sequencing. Doing so revealed that tumor progression correlates with increasing expression of genes involved in proliferation, interferon response, *IL2-STAT5*, and *TNFa*. These genes show temporal trends in our longitudinal single-cell transcriptomics data too. Cancer cells express decreasingly proliferation genes and increasingly IFN-response genes as the experiment progresses (Figure 2). (IL2-)stimulated Tregs peak at days 14-18 (Figure 4D–F). *TNFa* expression associates with the transcriptional heterogeneity of multiple immune cells types: TAMs (M1 archetype, peak at day 18, Figure 2), Group 1 ILCs (PC2, peak at days 14-18, Figure 4A), T helper cells (Th1 signature, peak at day 14, Figure 4C).

Similarly, expression of co-inhibitors (*CD274*, *IDO1*, *PDCD1*, *CTLA4*, *TIGIT*) and suppressive interleukins (*IL6*, *IL10*, *TGFB1*) were found to increase during the progression of human lung tumors[6]. In our longitudinal experiment, expression of co-inhibitors *Cd274*, *Pdcd1*, *Ctla4*, *Tigit* as well as suppressive interleukins Il10 and Tgb1 increased over time too, peaking at 18 (Figure S11B-C). *Ido1* and *Il6* were weakly expressed throughout the experiment or decreased over time (Figure S11B-C).

Thus, cancer and immune genes identified to peak at the most advanced stage of human lung tumors also increased over time in our longitudinal experiment but peaked at day 18, not at the latest time point (day 24). This difference could have multiple explanations. Ordering human tumors by histological stage may imperfectly reconstitute temporal dynamics. Differences in the size of host organisms could cause mouse — but not human — tumors to reach carrying capacity and thus limit tumor proliferation at the latest time point (day 24).

https://www.nature.com/articles/s41588-021-00911-1. Publisher: Nature Publishing Group.

[6] Céline Mascaux, Mihaela Angelova, Angela Vasaturo, Jennifer Beane, Kahkeshan Hijazi, Geraldine Anthoine, Bénédicte Buttard, Françoise Rothe, Karen Willard-Gallo, Annick Haller, Vincent Ninane, Arsène Burny, Jean Paul Sculier, Avi Spira, and Jérôme Galon. Immune evasion before tumour invasion in early lung squamous carcinogenesis. *Nature 2019 571:7766*, 571(7766):570–575, June 2019. ISSN 1476-4687. doi: 10.1038/s41586-019-1330-0. URL https://www.nature.com/articles/s41586-019-1330-0. Publisher: Nature Publishing Group.

[7] Victoria Sopik and Steven A. Narod. The relationship between tumour size, nodal status and distant metastases: on the origins of breast cancer. *Breast Cancer Research and Treatment*, 170(3):647–656, August 2018. ISSN 1573-7217. doi: 10.1007/s10549-018-4796-9. URL https://link.springer.com/article/10.1007/s10549-018-4796-9. Company: Springer Distributor: Springer Institution: Springer Label: Springer Number: 3 Publisher: Springer US.
